# Mucoricin binding to β-glucan sites on germinating Mucorales spores disrupts neutrophil swarming to promote pathogenicity

**DOI:** 10.1101/2025.10.28.685056

**Authors:** Stavroula Baimpa, Matthaios Sertedakis, Elias Drakos, Tonia Akoumianaki, Alexandra Vatikioti, Yiyou Gu, Sanam Dolati, Anastasios Koutsopoulos, Ashraf S. Ibrahim, Georgios Chamilos

**Author notes:** Lead Contact: Georgios Chamilos, Laboratory of Clinical Microbiology and Microbial Pathogenesis, School of Medicine, University of Crete, Unit 3D72, Stavrakia, Voutes, Heraklion, Crete, Greece, 71110.

## Abstract

Mucormycosis is an emerging, life-threatening infection caused by Mucorales fungi with incompletely understood immunopathogenesis. We found that unlike other fungal pathogens, Mucorales trigger coordinated neutrophil clustering (swarming) upon swelling of fungal spores. Neutrophil swarms efficiently eliminate germinating Mucorales via ROS production, upon sensing of β-glucan by complement receptor 3 (CR3). Impaired neutrophil clustering in mucormycosis-predisposing conditions, allows for mucoricin toxin production by germinating Mucorales. Released mucoricin selectively binds to β-glucan on the surface of Mucorales, creating a molecular trap that induces neutrophil apoptosis and swarming disruption, leading to invasive fungal disease. Suppression of mucoricin, either genetically or physiologically by albumin-bound free fatty acids, stabilizes neutrophil clusters leading to abrogation of fungal pathogenicity. Additionally, prophylactic GM-CSF administration prevents mucoricin-induced neutrophil death, and protects mice from mucormycosis. Our work identifies neutrophil swarming as the primary target of mucoricin and provides compelling evidence for therapeutic harnessing of this pathway to improve mucormycosis outcome.

## Introduction

Mucorales fungi cause mucormycosis, an emerging, life-threatening disease of incompletely understood pathogenesis and limited therapeutic options^1–3^. *Rhizopus* species are responsible for the vast majority (∼70%) of mucormycosis cases worldwide^1,4^. Pulmonary mucormycosis, one of the most common manifestations of the disease, is associated with mortality rates that exceed 50% and approach 100% upon dissemination^1,2,5^. Extensive tissue necrosis, largely mediated by the mycotoxin mucoricin^6^, is a hallmark feature of mucormycosis that underlies the term “black fungus” and largely contributes to the poor disease outcome^1^. Understanding the early events in immunopathogenesis of mucormycosis is crucial for developing novel therapeutic strategies and improving disease outcome, which has been recently acknowledged as a high research priority by World Health Organization (WHO)^7^.

Similar to other invasive mold infections (IMIs), mucormycosis occurs in immunocompromised patients with profound defects in the numbers and/or function of neutrophils, as a result of chemotherapy-induced myelosuppression, transplantation, or treatment with high doses of corticosteroids^1^. However, unlike other IMIs, mucormycosis specifically affects an expanding group of patients with immunometabolic disorders, including diabetic ketoacidosis (DKA), increased iron availability, malnutrition, severe hypoalbuminemia^8^, critical illness, sepsis and COVID-19, through incompletely characterized mechanisms^1,2,4,5,9,10^. Importantly, host metabolites directly modulate the pathogenicity of Mucorales. Specifically, albumin-bound free fatty acids (FFAs) selectively inhibit the growth and pathogenicity of Mucorales^8^. On the other hand, excessive β-hydroxybutyrate (BHB) production in DKA promotes fungal growth and facilitates angioinvasion by upregulating the expression of fungal CotH invasins and their endothelial receptors^11^. The mechanisms by which metabolic abnormalities inhibit specialized immune responses against Mucorales, leading to development of mucormycosis, remain currently unknown.

Professional phagocytes, mainly alveolar macrophages (AMs) and recruited neutrophils, confer sterilizing immunity against Mucorales in the lung^12^. AMs constantly phagocytose and eliminate inhaled Mucorales spores^13^, while neutrophils comprise the main line of host defense against extracellular fungi that evade immune surveillance by AMs. In contrast to other airborne fungal pathogens^14^, germinating (swollen) *Rhizopus delemar* spores subvert all physiological host defense mechanisms, leading to acute lethality following pulmonary infection in immunocompetent mice^13^. A crucial gap of knowledge in immunopathogenesis of mucormycosis is the molecular understanding of Mucorales-neutrophil dynamic interplay during the early stages of infection.

Neutrophils combat invading pathogens through various immune effector mechanisms, including phagocytosis, reactive oxygen species (ROS) production, degranulation, NETosis, and swarming^15,16^. Swarming is characterized by the coordinated clustering of neutrophils in response to autocrine Leukotriene B4 (LTB4) release following sterile injury, or infections by parasites, bacterial aggregates or fungal hyphae^17–21^. Specialized neutrophil responses against large-size pathogens, including germinating fungi, mainly entail extracellular ROS production^22^, NETosis^23^, and swarming^19^. However, the outcome of neutrophil-fungal interplay is largely determined by the underlying disease context. Specifically, NETosis has a protective role against systemic, bloodstream *Aspergillus fumigatus* infection^24^, while it is dispensable for localized fungal infections^25^. Furthermore, while neutrophil swarming inhibits growth of fungal hyphae ex vivo^19^, excessive swarming promotes immunopathology in preclinical models of fungal pneumonia^26,27^. Therefore, the physiological role of neutrophil swarming in antifungal immunity is currently incompletely characterized.

In this study, we identify neutrophil swarming as a specialized host defense mechanism against Mucorales that is triggered by β-glucan exposure on germinating fungal spores. Furthermore, we uncover a novel pathogenetic mechanism that disrupts neutrophil swarming upon binding of released mucoricin to β-glucan sites on Mucorales cell wall. Under disease-predisposing conditions, mucoricin-β-glucan complex formation on germinating Mucorales spores acts as a molecular trap that rapidly induces apoptosis of recruited neutrophils and promotes invasive fungal disease. Of interest, inhibition of mucoricin expression by albumin-bound FFAs or abrogation of neutrophil apoptosis by prophylactic administration of granulocyte-macrophage colony-stimulating factor (GM-CSF) enhances neutrophil swarming and protects against mucormycosis. Overall, our study sheds light on molecular interactions during neutrophil-Mucorales interplay with direct translational impact for host-directed therapeutic strategies against mucormycosis.

## Results

### Neutrophil swarming is a specialized host defense mechanism against Mucorales

To explore the physiological interactions between Mucorales and neutrophils, we infected immunocompetent mice intratracheally with *R. delemar* spores and assessed the inflammatory immune response in the lung over time, by histopathology and immunohistochemistry for Periodic acid-Schiff’s (PAS) combined with myeloperoxidase (MPO). A robust neutrophil influx was detected in the lung within 3 h of infection, with evidence of organized neutrophil cell clusters surrounding extracellular spores of *R. delemar* (**Figure 1A, 1B**). This immunological response was suggestive of neutrophil swarming, which was confirmed by immunofluorescence staining of lung histopathology sections obtained from mice infected with fluorescence-labeled *R. delemar* spores (**Figure 1C**).

**Figure 1.**
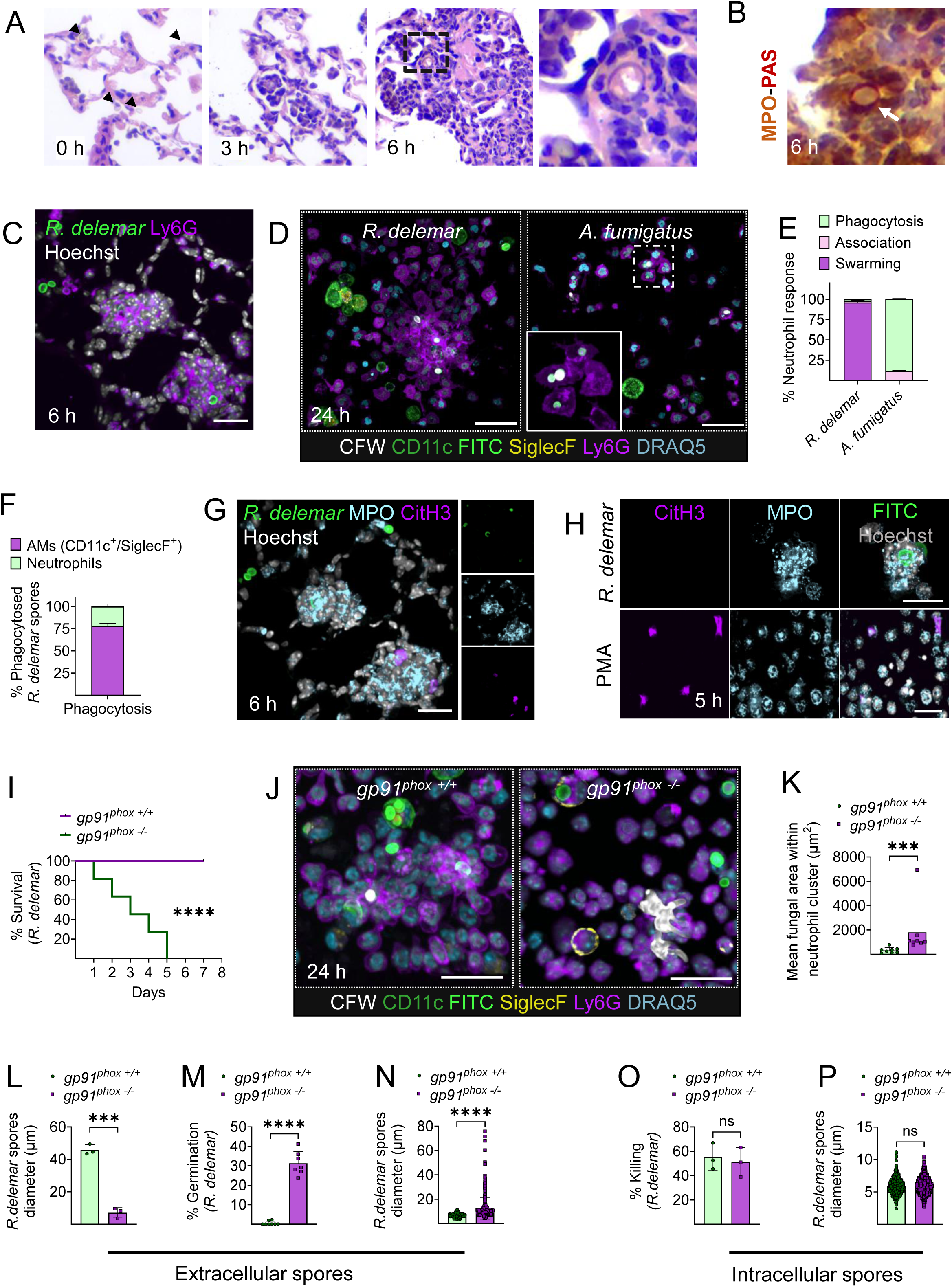
Neutrophil swarming is a specialized host defense mechanism against Mucorales. **(A)** Representative photomicrographs of the lungs from mice infected with dormant *R. delemar* spores (indicated with black arrows) and sacrificed 0, 3- and 6-hours (h) post infection. Lungs were counterstained with hematoxylin. **(B)** Representative photomicrograph of the lungs from mice infected with dormant *R. delemar* spores and sacrificed at 6 h of infection. Lungs were counterstained with PAS and MPO, indicating clearly the neutrophils in swarms around the spore (indicated with white arrow). **(C)** Representative confocal images of the lungs from mice infected as in (A) with FITC-labeled *R. delemar* and sacrificed at 6 h of infection. Lungs were stained for Ly6G and Hoechst. Scale bar, 20 μm. **(D)** Representative confocal images from BAL from immunocompetent mice at 24 h of infection with dormant *R. delemar* spores or *A. fumigatus* conidia. The fungal spores were prelabelled with FITC (green). BAL cells were allowed to attach in specialized, high-end microscopy plates (ibidi), and stained with Calcofluor White (CFW) to discriminate intracellular (FITC^+^/CFW^-^) vs extracellular spores (FITC^+^/ CFW^+^). Different immune cell subsets were stained with appropriate fluorescent-conjugated antibodies. Scale bar, 40 μm. **(E)** Data on quantification of different neutrophil responses in the lungs of immunocompetent mice infected with either *R. delemar* or *A. fumigatus* and evaluated by immunostaining of BAL at 24 h. **(F)** Data on quantification of the percentage of macrophages or neutrophils, that have phagocytosed *R. delemar* spores. **(G)** Representative confocal images of the lungs from mice infected with dormant *R. delemar* spores-prelabeled with FITC, and sacrificed at 6 h of infection. Lungs were stained by IHC for MPO and CitH3 to evaluate the degree of neutrophil extracellular trap (NET) formation. Scale bar, 20 μm. **(H)** Representative image from ex vivo infection of mouse bone marrow neutrophils, 4 h post infection. Scale bar, 30 μm. **(I)** Survival rates of immunocompetent *gp911^phox +/+^* (WT) mice and *gp91^phox -/-^* (CGD) mice (n=11 per group) infected with dormant *R. delemar* spores. The nonparametric log-rank test was used to determine differences in survival rates. **(J)** Immunofluorescence of BAL obtained 24 hours post infection from immunocompetent *gp91^phox +/+^* mice and *gp91^phox -/-^* mice, infected with dormant *R. delemar* spores. **(K)** Mean fungal area within neutrophil clusters in BAL of *gp91^phox +/+^* mice and *gp91^phox -/-^* mice, infected with dormant *R. delemar* spores (n= 2/group). \*\*\**P=*0.0003, Mann Whitney test. **(L)** Percentage of killed extracellular *R. delemar* spores 24 h post infection of immunocompetent *gp91^phox +/+^* and *gp91^phox -/-^* mice with dormant fungal spores. \*\*\**P*=0.0001, Unpaired t-test. **(M)** Representative data of the percentage of the extracellular germinating *R. delemar* spores 24 h post infection \*\*\*\**P* < 0.0001, Mann-Whitney test. **(N)** Representative data of the germination of extracellular *R. delemar* spores 24 h post infection. \*\*\*\**P* < 0.0001, Mann-Whitney test. **(O)** Percentage of killed *R. delemar* intracellular spores 24 h post infection of immunocompetent *gp91^phox +/+^* and *gp91^phox -/-^* mice with dormant fungal spores. Unpaired t-test. **(P)** Representative data of the germination of intracellular *R. delemar* spores 24 h post infection. *P*=0.1062, Mann-Whitney test.

Next, we performed comparative immunological analysis by confocal microscopy in bronchoalveolar lavage (BAL) cells obtained from mice infected with fluorescently labeled spores of *R. delemar* or *A. fumigatus* to evaluate the specificity of neutrophil swarming against Mucorales. Of interest, we found that the vast majority of extracellular *R. delemar* spores (≈85%) were contained within large neutrophil clusters (**Figure 1D, 1E, S1A**), while most intracellular spores were phagocytosed by macrophages (**Figure 1F, S1A**). In contrast, *A. fumigatus* spores were exclusively phagocytosed during interactions with neutrophils (**Figure 1D, 1E**). Furthermore, ex vivo studies in mouse bone-marrow derived neutrophils infected with either *A. fumigatus*, *R. delemar* or other Mucorales spp. confirmed that neutrophil swarming is a specialized immune response against Mucorales fungi (**Figure S1B-D**).

Considering that neutrophil swarming in response to other fungal pathogens is accompanied by the release of NETs within neutrophil clusters^19,20,27^, we evaluated NETosis during infection with *R. delemar*. Importantly, immunostaining for citrullinated histone H3, a specialized marker of NETosis, myeloperoxidase, and DNA revealed no evidence of NETosis within or around neutrophil swarms during *R. delemar* infection, both in vivo and ex vivo (**Figure 1G, 1H**).

To evaluate the role of neutrophil swarming in fungal clearance, we developed a protocol that allows simultaneous assessment of the viability of intracellular and extracellular fungal spores during pulmonary infection with *R. delemar*. Briefly, BAL fluid was stained with Calcofluor White (CFW) to discriminate extracellular spores^28^. BAL cells were lysed by sonication to release intracellular spores, which were then cultured in appropriate media, and the fungal killing was assessed by measurement of germination rates by confocal microscopy (**Figure S2A**). Of interest, killing of intracellular *R. delemar* spores, which are mainly phagocytosed by AMs^13^ (**Figure 1F**), started at 3 h of infection, whereas killing of extracellular spores started at 6 h, correlating with the kinetics of neutrophil swarming (**Figure S2B**). Within 48 h of infection, a significant and comparable amount of intra- and extra-cellular fungal spores were killed, which was further validated by CFU counts of total lung homogenates (**Figure S2B, S2C**). These findings identify neutrophil swarming as a central host defense mechanism against Mucorales.

We then sought to identify the effector mechanism(s) involved in the killing of *R*. *delemar* spores within neutrophil clusters. NADPH oxidase-mediated extracellular ROS production has been shown to inhibit ex vivo growth of *Candida* and *Aspergillus* hyphae during interactions within neutrophil swarms^19^. Furthermore, patients with chronic granulomatous disease (CGD), a primary immunodeficiency caused by genetic mutations in each of the subunits of NADPH oxidase, display increased susceptibility to mucormycosis^29^. Therefore, we infected intratracheally, CGD (*gp91^phox-/-^*) and control (*gp91^phox+/+^*) immunocompetent mice with *R. delemar* and assessed both survival and the immune response in the pulmonary space. Importantly, we found that infection with dormant *R. delemar* spores resulted in fulminant disease in *gp91^phox-/-^* mice, with mortality rates of 100% within five days (**Figure 1I**). Analysis of immune responses, following pulmonary infection of CGD and control mice with *R. delemar,* revealed that CGD neutrophils failed to restrict fungal growth within neutrophil clusters (**Figure 1J, 1K**), despite evidence of robust swarming. In contrast, there was no evidence of growth for *R. delemar* inside the AMs of either control or CGD mice (**Figure 1SD**). Accordingly, we found that killing of extracellular *R. delemar* spores in the lung was abolished in CGD mice (**Figure 1L-N**), whereas intracellular fungal killing by AMs was unaffected and comparable between CGD vs. control mice (**Figure 1O, 1P**). Collectively, these findings provide novel insights on the physiological mechanisms of killing of germinating Mucorales spores within neutrophil swarms via extracellular ROS release, which is in contrast to the ROS-independent mechanism of killing of phagocytosed spores inside AMs. These studies also illustrate that professional phagocytes employ distinct and specialized host defense mechanisms against Mucorales, both intracellularly and extracellularly^30^.

### β-glucan exposure on germinating spores triggers neutrophil swarming via activation of complement receptor 3 (CR3)

To gain insight into the dynamics of activation of neutrophil swarming induced by Mucorales, we performed 4D time-lapse microscopy during interaction of mouse bone marrow neutrophils with *R. delemar*. We found that clustering of neutrophils around *R. delemar* spores was induced at ≈ 2 h of infection (**Figure 2A, Suppl Video 1**) and was associated with a significant increase of spore diameter (swelling) (**Figure 2B**). To clarify whether swarming activation is induced by a secreted factor during the early phase of fungal germination, we stimulated neutrophils with paraformaldehyde (PFA)-inactivated *R. delemar* spores at different stages of growth (0 h, 2 h, 4 h). We found that, in contrast to PFA-inactivated dormant spores (0 h), PFA-inactivated *R. delemar* spores previously swollen for either 2 h or 4 h, triggered robust and comparable activation of neutrophil swarming with that induced by infection with live fungal spores (**Figure 2C, 2D**), which was also confirmed in experiments with human neutrophils (**Figure S3A, S3B**). These finding suggested that neutrophil swarming is activated upon the unmasking of an immunostimulatory molecule on cell wall of germinating *R. delemar* spores following melanin removal ^13,31,32^. Indeed, purified *R. delemar* melanin ghosts failed to activate neutrophil responses (**Figure S4A, S4B**), whereas PFA-inactivated dormant, melanin-deficient *R. delemar* spores, generated by pharmacological inhibition of melanin production with baathocuproinedisulfonic acid disodium salt (BCS)^13^, strongly activated neutrophil swarming (**Figure S4C**).

**Figure 2.**
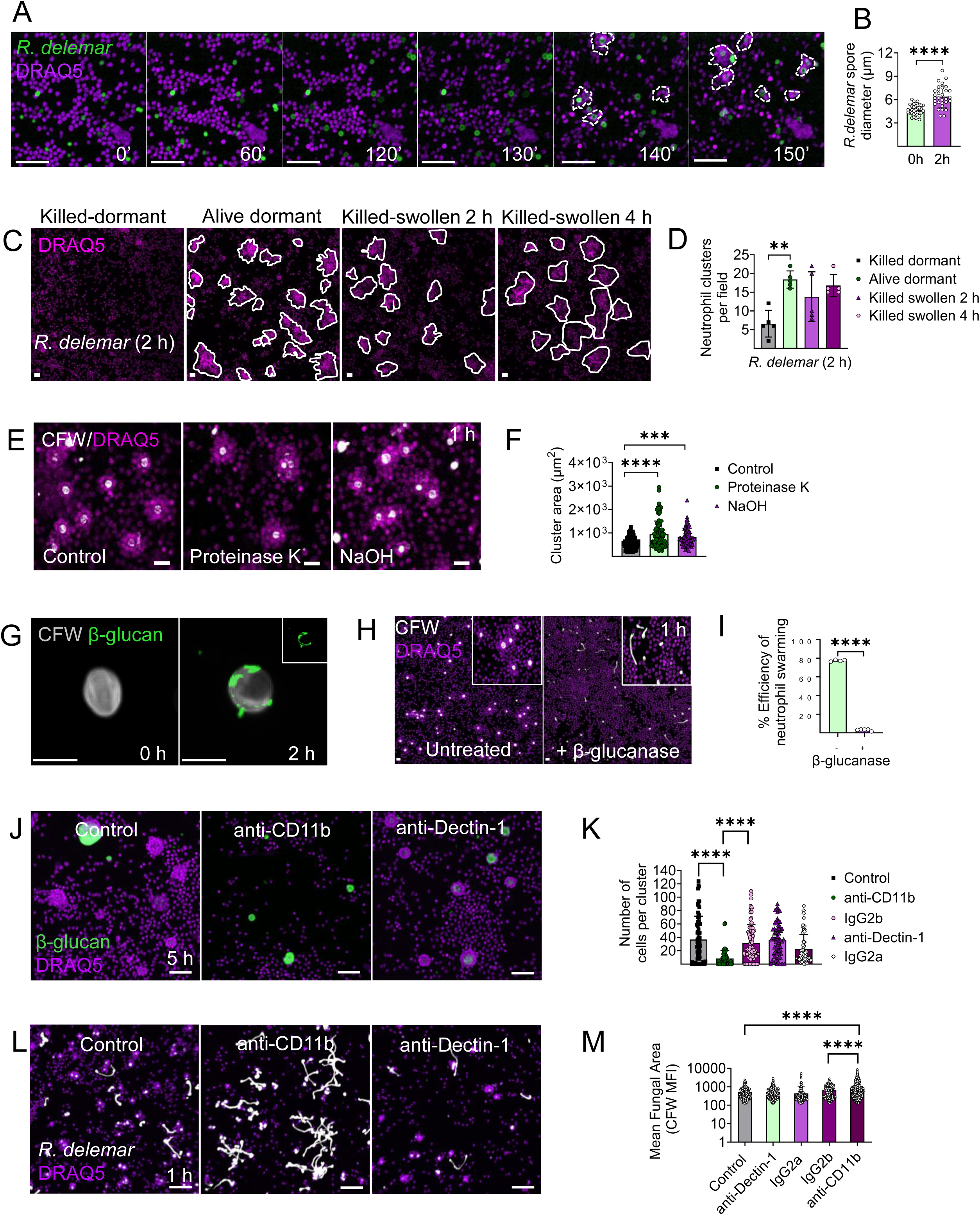
β-glucan exposure on the cell wall of germinating *R. delemar* spores triggers neutrophil swarming via activation of complement receptor 3 (CR3). **(A)** Representative time-lapse images of mouse bone marrow isolated neutrophils challenged with dormant *R. delemar* spores. Scale bar, 50 μm. **(B)** Data on quantification of spore diameter of dormant and swollen *R. delemar* spores. \*\*\*\**P* <0.0001, Mann-Whitney test. **(C)** Representative images of bone marrow neutrophil infected with alive dormant and PFA-killed dormant, 2- and 4 h swollen *R. delemar* spores. Scale bar, 50 μm. **(D)** Number of neutrophil clusters per field following infection with swollen PFA-killed *R. delemar*. \*\**P* =0.021, One-way ANOVA. **(E)** Representative images of bone marrow-isolated neutrophils, challenged with swollen *R. delemar* spores treated with Proteinase K and NaOH. Scale bar, 20 μm. **(F)** Cluster area formed by bone marrow isolated neutrophils, challenged with swollen *R. delemar* spores treated with Proteinase K and NaOH. \*\*\**P* < 0.0001, \*\*\*\**P* < 0.0001 One-way ANOVA. **(G)** Representative images of immunofluorescence staining of β-glucan at dormant stage and 2 h swollen *R. delemar* spores. Scale bar, 5 μm. **(H)** Representative images of murine bone-marrow derived neutrophils infected with 4h swollen β-glucanase treated *R. delemar* spores. Scale bar, 10 μm. **(I)** Data on neutrophil swarming efficiency against 4 h swollen β-glucanase treated *R. delemar* spores. *\*\*\*\*P <0.0001,* Unpaired t-test. **(J-K)** Effects of inhibition of CR3 or Dectin-1 on neutrophil swarming against large β-glucan particles. *\*\*\*\*P <0.0001*, One-way ANOVA. **(L-M)** Effects of CR3 or Dectin-1 blockade on neutrophil-mediated inhibition of germination of *R. delemar* swollen spores. *\*\*\*\*P <0.0001,* One-way ANOVA.

To identify the immunostimulatory molecule that activates neutrophil swarming, we systematically removed different fungal cell wall surface components from swollen *R. delemar* spores. We initially removed surface proteins from the cell wall of PFA-inactivated swollen *R. delemar* spores, either enzymatically following overnight incubation with Proteinase K, or chemically by treatment with NaOH. Of interest, we found that surface protein removal enhanced neutrophil clusters surrounding swollen fungal spores (**Figure 2E, 2F**). Since NaOH treatment removes alkaline-soluble polysaccharides^33^, we focused our studies on immunostimulatory alkali-insoluble polysaccharides. In previous work, we identified that β-glucan exposure during germination of Mucorales spores activates Dectin-1 mediated Th17 responses^34^. Of interest, immunostaining for Fc-Dectin-1^31^ demonstrated β-glucan exposure on cell wall surface of swollen *R. delemar* spores at 2 h of growth (**Figure 2G**). Furthermore, enzymatic digestion of β-glucan on swollen *R. delemar* spores completely abolished neutrophil swarming following stimulation of mouse bone marrow neutrophils (**Figure 2H, 2I**).

Because neutrophil swarming occurs in response to large-size pathogens or bioparticle aggregates^35^, we assessed whether the larger size of Mucorales spores than those of *A. fumigatus* accounts for selective activation of swarming. Therefore, we exposed dormant *A. fumigatus* conidia to olorofim, an inhibitor of dihydroorotate dehydrogenase that inhibits polarized growth, while permitting isotropic growth^36^, to generate swollen conidia with a size comparable to that of swollen *R. delemar* spores. We found that olorofim-treated *A. fumigatus* swollen conidia, at the size of *R. delemar* swollen spores (≈ 8 μm), activated robust neutrophil swarming and displayed a high level of β-glucan surface exposure (**Figure S5A, S5B, S5C**). These results illustrate a size-dependent mechanism of neutrophil swarming activation following β-glucan exposure on germinating fungal spores.

Next, we explored the molecular mechanism of neutrophil swarming activation by *R. delemar* β-glucan. Fungal β-glucan activates both Dectin-1 and complement receptor 3 (CR3) in neutrophils^15,25^. Nonetheless, purified β-glucan bioparticles activate neutrophil swarming via CR3, independently of Dectin-1 signaling^35^. Therefore, we inhibited either CR3 or Dectin-1 in mouse bone marrow neutrophils with the use of specific monoclonal antibodies and evaluated the immune response following stimulation with purified large β-glucan particles or infection with live *R. delemar* spores. Importantly, inhibition of CR3 with the use of a blocking Ab specific for the CD11b domain^37^, almost completely abolished neutrophil swarming in response to β-glucan particles (**Figure 2J, 2K**) or germinating *R. delemar* spores (**Figure 2L, 2M**). In contrast, inhibition of Dectin-1 receptor had no apparent effect on neutrophil clustering against β-glucan particles (**Figure 2J, 2K**). Furthermore, inhibition of CR3-mediated neutrophil swarming resulted in unrestricted growth of *R. delemar* as compared to treatment with either Dectin-1 inhibitory antibody or the corresponding isotype control Abs (**Figure 2L, 2M**). These findings demonstrate that neutrophil swarming against *R. delemar* spores is mediated by CR3 and activation of this signaling pathway is crucial for the restriction of fungal growth.

### Germinating Mucorales spores induce neutrophil death through direct contact and disrupt neutrophil swarming

Germinating (swollen) Mucorales spores induce profound necrosis of lung tissue and rapid death of immunocompetent mice, via incompletely characterized host evading mechanisms^13^. Given that neutrophils comprise the main line of host defense against germinating Mucorales spores, we investigated the dynamics of neutrophils-Mucorales interplay following pulmonary infection with dormant vs swollen *R. delemar* spores. Importantly, we found that in contrast to dormant *R. delemar* spores, those pre-swollen in medium for 4 h induced massive neutrophil death, disrupted neutrophil swarming and caused invasive disease within 6 h of pulmonary infection (**Figure 3A, 3B**).

**Figure 3.**
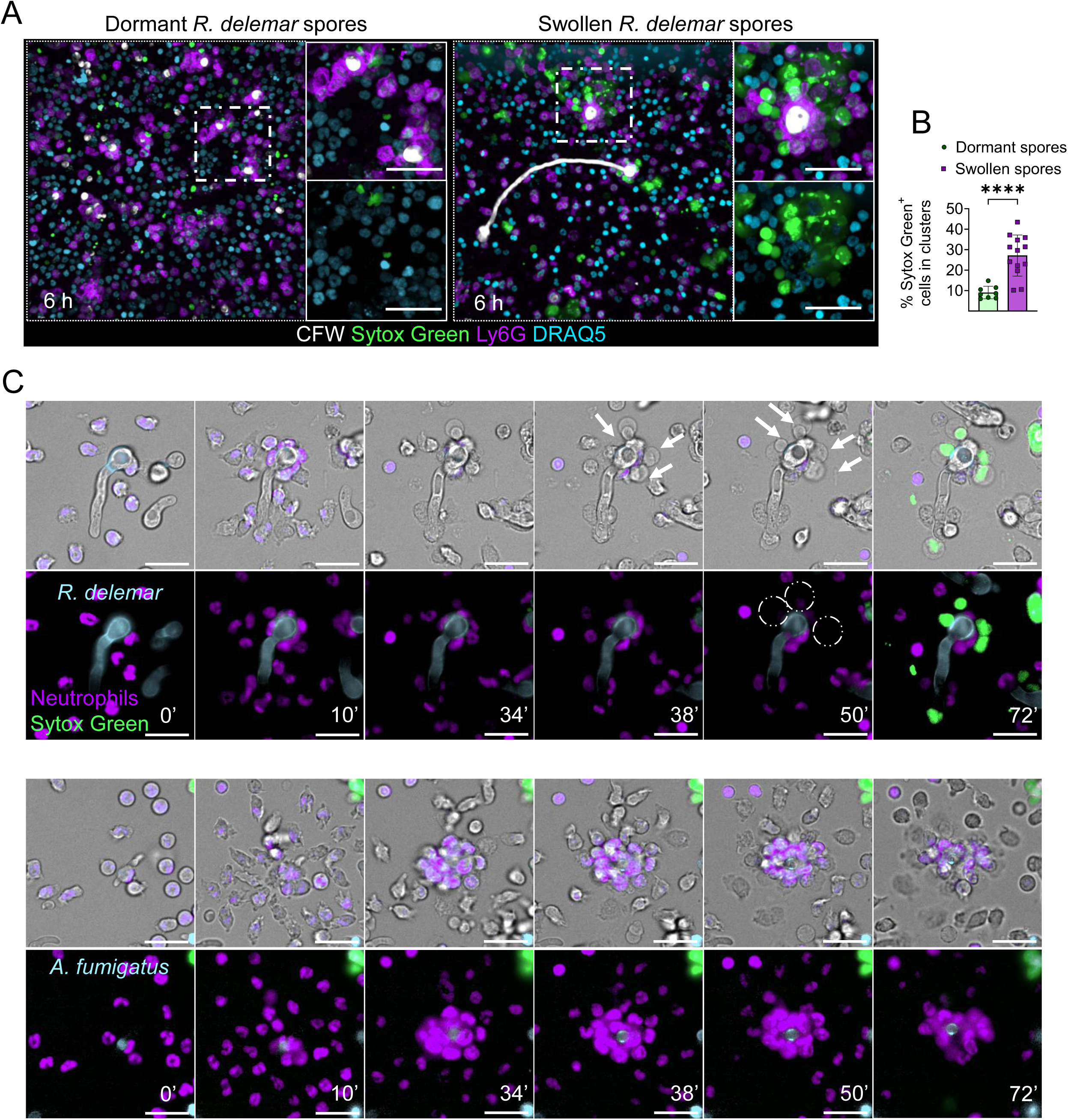
Germinating *R. delemar* spores disrupt neutrophil swarming via the induction of cell death upon direct contact. **(A)** Representative images of BAL cells isolated from mice infected with dormant and swollen *R. delemar spores* at 6 h post infection. Cell death is indicated by Sytox Green staining. Scale bar, 30 μm. **(B)** Percentage of Sytox Green^+^ neutrophils in clusters isolated from mice infected with dormant spores compared to the mice infected with swollen spores (n= 2/group). **** *P<0.0001*, Unpaired t-test. **(C)** Comparative live imaging studies of isolated mouse bone marrow neutrophils interacting with live germinating cells of *R. delemar* and *A. fumigatus.* Scale bar, 20 μm.

To elucidate the mechanism of neutrophil cell death induced by germinating/swollen *R. delemar*, we performed time lapse imaging studies of bone marrow-derived neutrophils infected with germinating spores of either *R. delemar* or *A. fumigatus*. We observed that during swarming, the attachment of neutrophils to germinating *R. delemar* spores triggered rapid host cell death within 30 minutes of infection, as evidenced by plasma membrane blebbing and nuclear condensation followed by staining with the cell impermeable nucleic acid dye Sytox Green (**Figure 3C, Suppl Video 2**). In contrast, neutrophils forming clusters around germinating *A. fumigatus* spores remained viable for prolonged periods (**Figure 3C**). Notably, germinating *R. delemar* spores pre-swollen in medium for 2 h strongly activated swarming without the induction of neutrophil death and were completely non-pathogenic in vivo (**Figure S6**). These data suggest that a cell wall-associated virulence factor exposed at ≈ 4 h of germination of *R. delemar* spores induces neutrophil death.

### Mucoricin binding to β-glucan sites on germinating *R. delemar* spores triggers neutrophil apoptosis

In view of the pivotal role of mucoricin in tissue necrosis following pulmonary infection with swollen *R. delemar* spores^8^, and considering that this hyphae-associated mycotoxin is highly expressed around 4-5 h of germination^6,8^, we explored the contribution of mucoricin to neutrophil death. We initially performed 4D live-cell imaging to analyze neutrophil responses against swollen spores of control (empty RNAi) vs mucoricin RNAi *R. delemar* strain, or large β-glucan particles. We found that in contrast to infection with control (empty RNAi), which resulted in disruption of neutrophil swarms over time, mucoricin-RNAi triggered the formation of stable and larger neutrophil clusters at comparable rates to those induced by β-glucan particles (**Figure 4A, S7 and Suppl Video 3**). Furthermore, inhibition of mucoricin activity by pre-incubation of swollen control (empty RNAi) *R. delemar* spores with galactose^6^, preserved neutrophil cluster integrity (**Figure S8Α**). These experiments identify mucoricin as the molecule in germinating *R. delemar* spores that disrupts neutrophil swarming.

**Figure 4.**
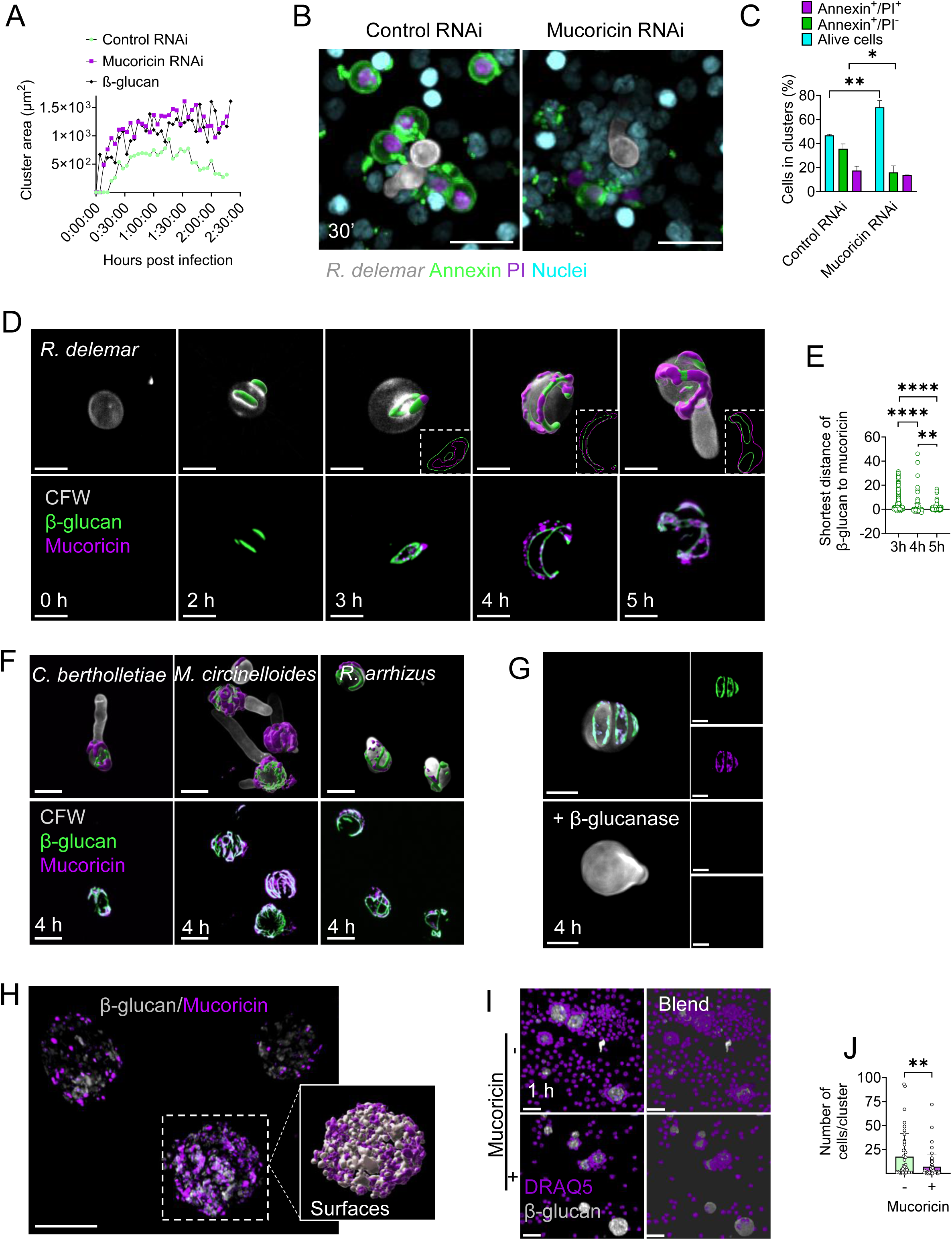
Binding of released mucoricin on β-glucan sites of the fungal cell wall triggers neutrophils death upon direct contact with germinating *R. delemar*. **(A)** Representative experiment on quantification of neutrophil swarming dynamics (cluster area of neutrophil nuclei over time) in time lapse imaging studies with mouse bone marrow neutrophils challenged with live *R. delemar* spores of control RNAi, mucoricin RNAi strains, or large β-glucan particles. **(B)** Representative images on assesment of apoptosis in mouse bone-marrow derived neutrophils challenged with control RNAi or mucoricin RNAi swollen *R. delemar* spores. Annexin-PI staining was performed at 30 min of infection. Scale bar, 20 μm. **(C)** Data on quantification of Annexin-PI staining of neutrophils in the experiment performed in B. *\*\*P =0.0051, *P =0.0148, two-way* ANOVA. **(D)** Representative images of immunofluorescence staining for β-glucan and mucoricin in different stages of growth of *R. delemar* spores following in vitro culture. Scale bar, 5 μm. **(E)** Quantification of shorter distance of β-glucan to Mucoricin, indicating the relative spatial positioning of the one in relation to the other. *\*\*\*\*P <0.0001*, *\*\*P = 0.0047*, One-way ANOVA. **(F)** Representative images of immunofluorescence staining for β-glucan and mucoricin in 4 h of growth of *C. bertholletiae*, *M. circinelloides* and *R. arrhizus*. Scale bar 10μm. **(G)** Representative image of 4 h swollen *R. delemar* treated or not with β-glucanase, stained for β-glucan and mucoricin. Scale bar, 5 μm. **(H)** Representative images of immunofluorescence staining of β-glucan particles incubated or not with purified mucoricin toxin. Scale bar, 20 μm. **(I)** Representative images of isolated mouse bone marrow neutrophils interacting with β-glucan particles incubated or not with purified mucoricin toxin. Scale bar, 30 μm. **(J)** Representative counting of the number of neutrophils in clusters around with β-glucan particles incubated or not with purified mucoricin toxin. \*\**P=0.0095*, Unpaired t-test.

Next, we performed immunostaining for Annexin V and Propidium Iodide (PI) at early time points of neutrophil infection with swollen spores of mucoricin or control RNAi *R. delemar* strains^6^ to explore the mechanism of mycotoxin-induced neutrophil death. Of interest, we found evidence of massive cell death (Annexin V^+^/PI^+^) in neutrophil clusters at 30 min, almost exclusively following infection with swollen spores of the empty RNAi strain (**Figure 4B, 4C**). This result suggests that mucoricin expression on Mucorales cell wall surface induces neutrophil apoptosis upon direct contact with germinating spores.

To gain insight on the spatial and temporal dynamics of neutrophil interaction with fungal cells we performed simultaneous immunostaining for mucoricin and β-glucan during different stages of growth of *R. delemar* spores. Of interest, mucoricin expression on swollen spores was delayed, when compared to β-glucan. Specifically, β-glucan was exposed on the surface of swollen spores within 2 h of growth (**Figure 4D**). On the contrary, mucoricin was detected at 3 h of growth underneath the fungal polysaccharide layer and became exposed on the outer surface of swollen spores and accessible to immune cells only after 4 h of growth (**Figure 4D, 4E**). Notably, immunostaining and 3D surface modeling revealed a striking co-localization of mucoricin and β-glucan with the former overlaying the later after 4 h of growth of *R. delemar* spores (**Figure 4D, 4E**); this pattern of mucoricin-β-glucan association was also observed in other Mucorales species (**Figure 4F**). These findings suggested the selective binding of released mucoricin on β-glucan sites of Mucorales cell wall. To test this hypothesis, we performed β-glucan removal on 4 h swollen spores by enzymatic digestion and assessed the effect on mucoricin expression. Importantly, β-glucanase treatment resulted in the simultaneous removal of β-glucan and surface-bound mucoricin, confirming their direct association (**Figure 4G**).

In view of the well-characterized mycotoxin-binding properties of β-glucan^38–40^, we assessed the binding affinity of purified mucoricin to β-glucan particles and evaluated the functional outcomes. Immunostaining studies revealed that mucoricin effectively bound on the surface of β-glucan particles (**Figure 4H**). Of interest, we found a significant reduction in the number of neutrophil cells in clusters following stimulation with mucoricin-coated β-glucan particles in comparison to control uncoated β-glucan particles (**Figure 4I, 4J**). Because mucoricin release during hyphal growth induces cell death of epithelial and endothelial cells^6^, we evaluated the effect of soluble mucoricin on neutrophil swarming. Purified mucoricin added in culture media resulted in disruption of pre-formed neutrophil clusters surrounding β-glucan particles (**Figure S8B, S8C**). Notably, neutrophils were significantly more susceptible to mucoricin-mediated cytotoxicity than epithelial and endothelial cells (**Figure S8D**). Collectively, these studies reveal a novel virulence strategy employed during the early stage of Mucorales growth, in which immunostimulatory β-glucan on the fungal cell wall surface acts as a “bait” to recruit neutrophils, while the binding of released mucoricin on immune activating sites induces neutrophil death and facilitates immune evasion and invasive hyphal growth.

We have recently discovered that albumin-bound physiological FFAs abrogate Mucorales pathogenicity in vivo by inhibiting mucoricin expression^8^. We hypothesized that inhibition of mucoricin release by albumin-bound FFAs will result in restoration of neutrophil swarming against swollen Mucorales spores and fungal clearance. We initially infected mice with swollen *R. delemar* spores with or without pre-exposure to albumin-bound FFAs and assessed neutrophil responses in BAL fluid at 6 h of infection. Importantly, we found that pre-incubation of Mucorales spores with albumin-bound FFAs triggered robust neutrophil swarming around fungal spores and abrogated neutrophil cell death (**Figure S9A-C**) These results were further validated in ex vivo experiments (**Figure S9D, S9E**). Overall, these studies uncover a physiological metabolic host defense mechanism that restores neutrophil swarming by inhibiting mucoricin-induced neutrophil death.

### Impaired neutrophil swarming in DKA promotes fungal pathogenicity and development of mucormycosis

Defects in neutrophil number, function, or chemotaxis are classical predisposing factors for invasive mold infections, including mucormycosis^1,12^. However, DKA and other forms of acidosis specifically predispose to mucormycosis via uncharacterized mechanisms of neutrophil dysfunction. Therefore, we investigated the effect of acidosis on swarming against β-glucan particles across a range of pH in standard culture media (**Figure 5A**). We found a significant impairment of neutrophil swarming at pH values below 7.2 (**Figure 5B, 5C**). Furthermore, physiological concentrations of BHB, the main ketone body produced in DKA, almost completely inhibited neutrophil swarming in a pH dependent fashion. Notably, reversal of acidosis following the addition of sodium bicarbonate to the medium restored neutrophil swarming (**Figure 5D, 5E**).

**Figure 5.**
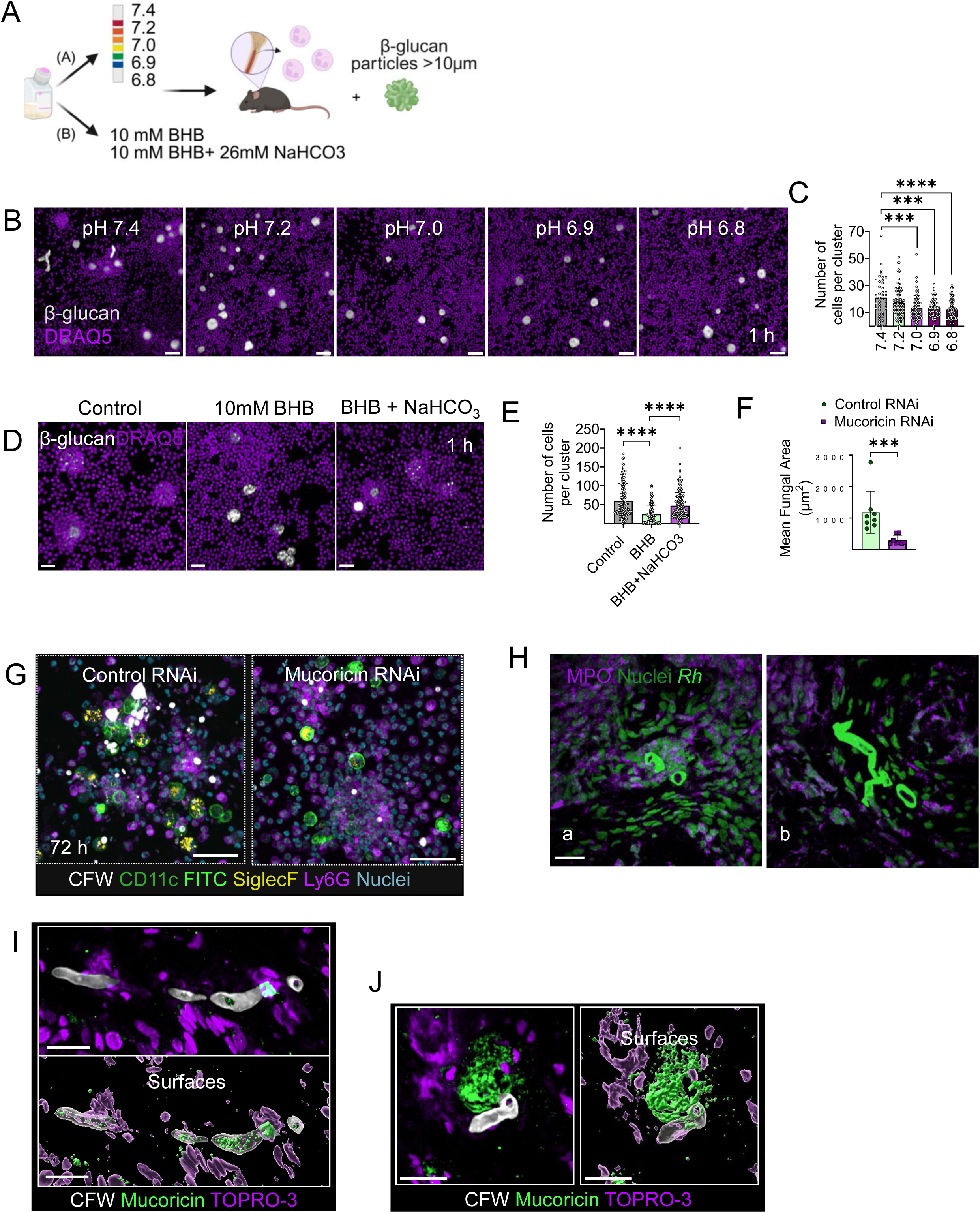
Acidosis inhibits neutrophil swarming and promotes the development of mucormycosis via mucoricin-induced cell death. **(A)** Schematic representation of the ex vivo experiments of bone-marrow isolated neutrophils, challenged with β-glucan particles (>10 μm) in medium with different pH. **(B)** Representative images of bone marrow-isolated neutrophils challenged with β-glucan particles in different pH. Scale bar, 50 μm. **(C)** Number of neutrophils in clusters around β-glucan particles in media with different pH. \*\*\*\**P*< 0.0001 One-way ANOVA. **(D)** Representative images of bone marrow-isolated neutrophils challenged with β-glucan particles in plain medium and medium supplemented with BHB or BHB+NaHCO3. Scale bar, 50 μm. **(E)** Number of neutrophils in clusters around β-glucan particles in plain medium and medium supplemented with BHB or BHB+NaHCO3. \**P*=0.0190, \*\**P*=0.0011, One-way ANOVA. **(F)** The mean fungal area of germinating Control RNAi or mucoricin RNAi *R. delemar* spores in BAL obtained from DKA mice, 72 hours post infection (n=2/group). \*\*\**P*=0.0002, Mann-Whitney test. **(G)** Representative confocal images from BAL obtained from DKA mice at 72 h of infection with Control RNAi or mucoricin RNAi dormant *R. delemar* spores. Scale bar, 50 μm**. (H)** Representative confocal images of histopathology section of tissue biopsy obtained from a DKA patient with rhino cerebral mucormycosis stained for fungal cells (CW), nuclei (TOPRO-3), and MPO^+^ cells (neutrophils). Scale bar, 20 μm. **(I-J)** Representative confocal images of histopathology section of tissue biopsy obtained from a DKA patient with rhino cerebral mucormycosis stained for fungal cells (CW), nuclei (TOPRO-3), and mucoricin. Scale bar, 20 μm.

We additionally evaluated the physiological relevance of mucoricin release on neutrophil responses in the lung of DKA mice^6^. Specifically, we infected DKA mice intratracheally with dormant spores of mucoricin RNAi or control (empty RNAi) *R. delemar* and analyzed immune responses in the BAL fluid at 72 h of infection. We observed that infection with control (empty RNAi) *R. delemar* spores promoted fungal growth within neutrophil clusters as a result of impaired neutrophil swarming and neutrophil death. In striking contrast, extracellular mucoricin RNAi *R. delemar* spores were contained within large neutrophil clusters without evidence of fungal germination or neutrophil death (**Figure 5F, 5G**). Collectively, these findings highlight the critical role of mucoricin in promoting early fungal invasion in DKA and other mucormycosis-predisposing conditions characterized by impaired activation of neutrophil swarming.

To further explore the physiological importance of swarming in mucormycosis we performed immunostaining for neutrophils in tissue biopsies obtained from a DKA patient with mucormycosis. Notably, we observed fungal outgrowth within areas of dissociated neutrophil clusters (**Figure 5Ha**). Additionally, extensive hyphal growth was associated with profound tissue necrosis in the absence of neutrophil clustering (**Figure 5Hb**). Immunostaining for mucoricin revealed that neutrophil cluster stability was disrupted both by direct engagement with fungal cell wall–bound mucoricin (**Figure 5I**) and by exposure to mucoricin released into adjacent tissue (**Figure 5J**). These findings further validate the pathogenetic role of neutrophil swarming disruption mediated by released mucoricin in development of mucormycosis.

### Inhibition of neutrophil apoptosis via prophylactic administration of GM-CSF restores neutrophil swarming and protects mice from mucormycosis

Adjunctive cytokine immunotherapies with GM-CSF, IFN-γ, and G-CSF, alone or in combination, have shown promising activity in preclinical studies and refractory cases of mucormycosis^41–46^. However, a systematic evaluation of the mechanism by which immunotherapy modulates the immune response against Mucorales has not yet been performed. Therefore, we explored the effect of individual cytokines on neutrophil responses against Mucorales, both ex vivo and in vivo. We initially preincubated murine bone marrow neutrophils with or without increasing concentrations of GM-CSF, IFN-γ, or G-CSF, infected them with swollen *R. delemar* spores and assessed the effects on neutrophil swarming and fungal growth at 8 h of infection. Importantly, we found that among all tested cytokines, GM-CSF significantly enhanced neutrophil cluster formation around *R. delemar* spores and effectively inhibited fungal germination ex vivo (**Figure 6A, 6B**).

**Figure 6.**
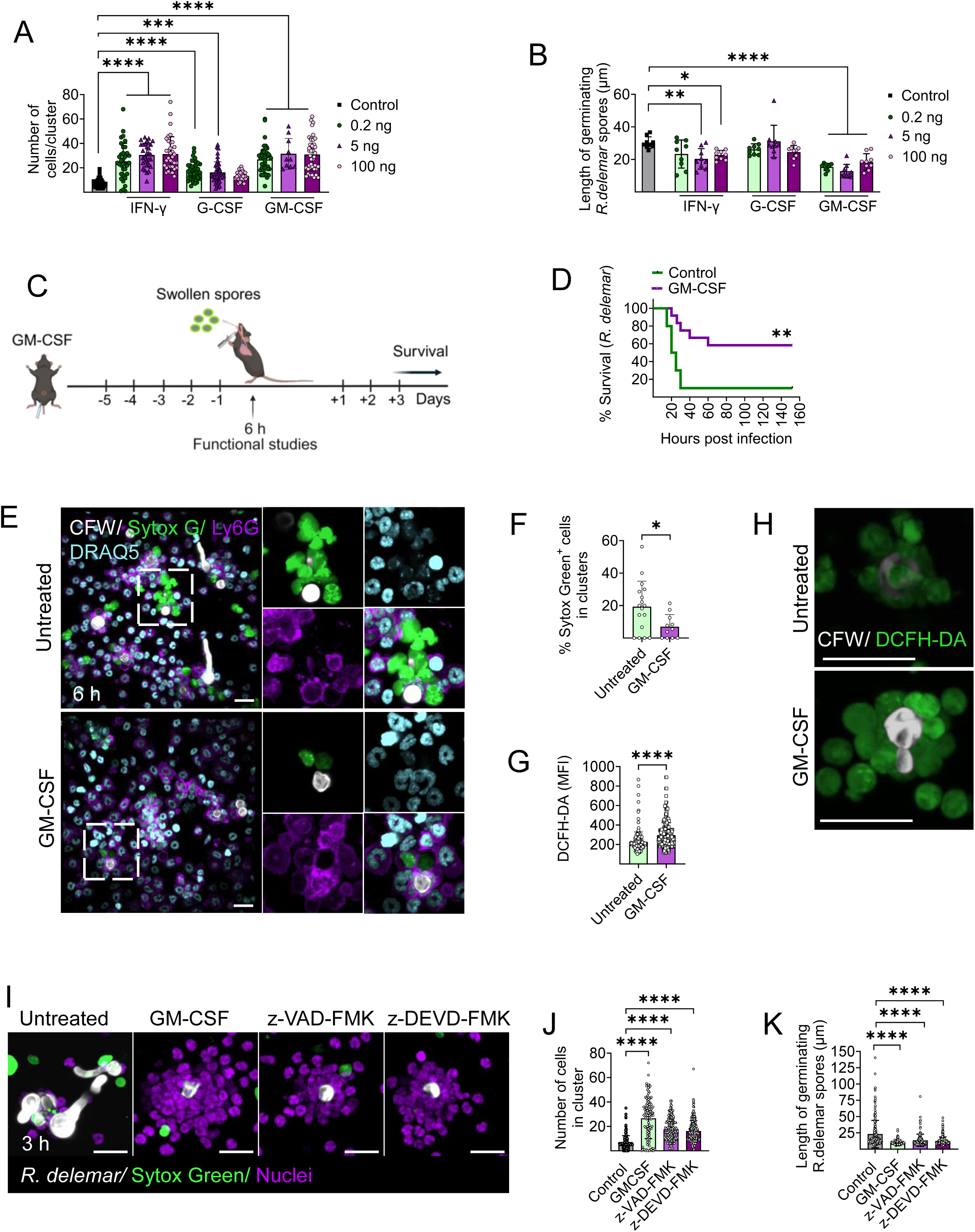
GM-CSF prevents mucoricin-induced apoptotic death of neutrophils and protects mice from mucormycosis via enhancing neutrophil swarming. **(A)** Quantification of the number of cells around *R. delemar* spores in the presence of different concentrations of IFN-γ, G-CSF and GM-CSF. \*\*\*\**P*<0.0001, \*\*\**P*=0.0002, \*\**P*=0.0012, One-way ANOVA. **(B)** The length of germinating *R. delemar* spores in the presence of different concentrations of IFN-γ, G-CSF and GM-CSF. \*\*\*\**P*<0.0001, \*\*\**P*=0.0002, \*\**P*=0.0022, \**P*=0.0308, One-way ANOVA. **(C)** Schematic representation of the prophylactic treatment regimen in mice using GM-CSF. **(D)** Survival of immunocompetent mice following infection with swollen spores of *R. delemar* with (n = 12) and without (n = 10) prophylactic administration of GM-CSF. The nonparametric log-rank test was used to determine differences in survival times. **(Ε)** Representative images obtained from BAL obtained 6 h post infection of mice in experiment performed in C. scale bar 30 μm. **(F)** Percentage of Sytox Green+ neutrophils in experiment performed in (C) *P=0.0222*, Unpaired t-test. **(G)** Measurement of ROS production in neutrophil clusters in BAL obtained 6 hours post infection of mice in experiment performed in (C). **** *P*< 0.0001, Mann-Whitney test. **(H)** Representative images obtained from BAL obtained 6 h post infection of mice in experiment performed in (C), indicating increased ROS production in clusters. scale bar 30 μm **(I)** Representative images showing the number of the cells in the clusters around *R. delemar* spores in the presence of GM-CSF, z VAD-FMK and z-DEVD-FMK. scale bar 20 μm **(J)** Quantification of the number of cells around *R. delemar* spores in the presence of GM-CSF, z VAD-FMK and z-DEVD-FMK **** *P*< 0.0001, One-way ANOVA. **(J)** The length of germinating *R. delemar* spores in the presence of GM-CSF, z-VAD-FMK and z-DEVD-FMK. **** *P*< 0.0001, One-way ANOVA.

Accordingly, we evaluated the protective effect of GM-CSF in the mouse model of pulmonary mucormycosis^8,13^. Immunocompetent mice received daily prophylactic administration of GM-CSF (**Figure 6C**) and then infected intratracheally with swollen *R. delemar* spores. Importantly, prophylactic treatment with GM-CSF resulted in significantly higher survival rates compared to control untreated mice (**Figure 6D**). Furthermore, analysis of immune responses in BAL cells at 6 h post-infection revealed that lung-infiltrating neutrophils remained viable and formed large clusters that effectively contained swollen *R. delemar* spores (**Figure 6E**). In contrast, BAL samples from untreated mice displayed excessive neutrophil death, swarming disruption and invasive hyphal growth within neutrophil clusters (**Figure 6E, 6F**). Lung-infiltrating neutrophils from GM-CSF-treated mice also produced significantly higher levels of ROS within the swarms as compared to those from untreated neutrophils (**Figure 6G, 6H**). Finally, we evaluated the role of anti-apoptotic properties of GM-CSF^47^ in enhancement of neutrophil swarming against Mucorales, by comparative studies with the selective inhibitor of caspase-3, z-DEVD^48^, and the pan-caspase inhibitor, z-VAD-FMK^49^. Notably, the immune augmenting effects of GM-CSF on neutrophil swarming against Mucorales were also apparent following pre-treatment with apoptosis inhibitors (**Figure 6I-6K**). These findings suggest that GM-CSF promotes both neutrophil survival and effector function and holds promise as an adjunctive immunotherapy for mucormycosis.

## Discussion

Dissecting early events in immunopathogenesis of mucormycosis is an unmet need to design better therapies and improve disease outcome. We initially characterized physiological immune responses in the lung of immunocompetent mice during infection with Mucorales in comparison to *A. fumigatus.* We found that Mucorales spores that had not been phagocytosed by AMs triggered robust neutrophil swarming. In contrast, neutrophils phagocytosed all encountered *A. fumigatus* conidia. Previous studies demonstrate a controversial role of neutrophil swarming in antifungal immunity. Specifically, concordant activation of NETosis in neutrophil swarms drives immunopathology in *Pneumocystis jirovecii* pneumonia^50^. Similarly, neutrophil clustering mediates pulmonary vascular occlusion during intravascular sepsis by *Candida*^20^. Finally, excessive neutrophil clustering due to unabated LTB4 release results in inflammatory deregulation during respiratory infection by *A. fumigatus* in CGD mice^51^.

Our study identified swarming as a specialized neutrophil response against Mucorales. Furthermore, experiments in control and CGD mice demonstrated that NADPH oxidase-dependent ROS production within neutrophil clusters is the primary mechanism of killing of germinating Mucorales spores in the lung. These findings are consistent with a NADPH oxidase dependent mechanism of restriction of *Candida* and *Aspergillus* hyphae by neutrophil swarming ex vivo^15^. Our work reveals the critical role of neutrophil swarming in host defense against Mucorales and offers mechanistic insights on the increased susceptibility of CGD patients to mucormycosis^29,52^.

Our experiments into the molecular mechanism of Mucorales activation of neutrophil swarming point to β-glucan exposure during the early germination phase (swelling) of Mucorales spores as a major driver of events. First, we confirmed the immune-silencing role of melanin on the surface of dormant Mucorales spores through β-glucan masking^13^. Second, pharmacological modulation of *A. fumigatus* spore size further demonstrated that the larger size of Mucorales spores drives neutrophil swarming in response to β-glucan exposure. These findings are in line with previous studies on the size-dependent activation of neutrophil swarming against aggregates of fungal bioparticles^19^ and offer a mechanistic explanation for the specificity of this host defense mechanism against Mucorales. Finally, blocking experiments with specific inhibitory antibodies against the major β-glucan pattern recognition receptor (PRR) in neutrophils, Dectin-1 and CR3^25^, identified CR3 as the main PRR that triggers neutrophil swarming against Mucorales. Notably, CR3 cross talk with VLA-3 vs VLA-5 integrin receptors, which depends on the composition of extracellular matrix, regulates activation of neutrophil swarming or NETosis in response to β-glucan, respectively^53^. A recently identified inhibitory C-type lectin receptor, MICL, blocks NETosis upon sensing of DNA^24^. Future studies should investigate whether differences in cell wall composition of other fungi when compared to Mucorales could account for the reported activation of NETosis by β-glucan in neutrophil clusters^54^.

To gain insight on the dynamic interaction of neutrophils with germinating Mucorales, we analyzed immune responses in the lung of immunocompetent mice following infection with fungal spores at different stages of growth. Strikingly, we found that germinating (swollen) Mucorales spores induced massive neutrophil death and swarming disruption, leading to fulminant infection in immunocompetent mice. Time lapse imaging studies revealed an apoptotic mechanism of neutrophil death within minutes of neutrophil contact with the cell wall of Mucorales. Further experiments with the use of mucoricin RNAi strains provided unambiguous evidence for the role of this novel mycotoxin^6^ in the disruption of neutrophil swarming. Of interest, neutrophils are resistant to the cytotoxic effects of mycotoxins^55^, in comparison to other phagocytes. Accordingly, neutrophil death has not been observed during physiological interactions with germinating forms of other human fungal pathogens^14^. Therefore, our study identified a previously uncharacterized virulence property of mucoricin to directly attack the immune system and promote lethal disease.

Our findings on mucoricin-induced neutrophil apoptosis upon direct contact with swollen Mucorales spores suggested for mycotoxin binding on the fungal cell wall surface. Advanced microscopy imaging studies revealed a striking colocalization of mucoricin with sites of β-glucan exposure on Mucorales spores. Enzymatic digestion of β-glucan along with functional studies following incubation of purified mucoricin with β-glucan particles confirmed that β-glucan-bound mucoricin triggered neutrophil death. The toxin-binding properties of β-glucan have been explored in industrial applications for absorption of fungal mycotoxins^38–40,56^. Notably, our study is the first to demonstrate the ability of a microbial toxin to bind on immunostimulatory cell wall polysaccharides, creating a molecular trap that attacks recruited immune cells. Mucoricin binding on fungal cell wall β-glucan could be the result of evolutionarily pressure during interaction with other environmental organisms or predators. In particular, *Saccharomyces* released K2 killer mycotoxin binding on cell wall β-1,6-glucan provides a growth advantage during competition with other fungal cells^57,58^. Furthermore, toxin production by endosymbiotic bacteria of *Rhizopus microspores* protects fungal cells from micropredators, such as frugivorous amoebas and nematodes^59^. It is tempting to speculate that the swarming behavior of frugivorous amoebae during their environmental interactions with toxin-producing fungi^60,61^ may have served as an evolutionary precursor of neutrophil swarming.

We have recently identified albumin-bound FFAs as physiological metabolic host effectors that inhibit Mucorales pathogenicity and mucoricin release^8^. In this study, we discover that the FFA-mediated inhibition of mucoricin release restores neutrophil swarming to prevent development of mucormycosis. These findings uncover a unique connection between metabolic serum host defense and cellular innate immunity against an emerging fungal pathogen and provide further evidence on the central role of neutrophil swarming against mucormycosis.

We further explored the physiological importance of mucoricin-mediated neutrophil cell death in the DKA model of mucormycosis. Of interest, we initially identified that acidosis, similar to other predisposing conditions for mucormycosis^62^, abolished activation of neutrophil swarming, in a pH dependent way. This finding is consistent with inhibition of LTB4 release by neutrophils during acidosis as a result of depolarization of cellular membranes^63^. Furthermore, we discovered that in the DKA mouse model of mucormycosis, impaired neutrophil clustering around fungal spores allowed for mucoricin-mediated swarming disruption by germinating hyphae of Mucorales. In contrast, genetic ablation of mucoricin resulted in stable neutrophil cluster formation around Mucorales spores and inhibition of hyphal growth. These findings were also validated in studies using human histopathology biopsies from DKA patients with mucormycosis. Collectively, these studies illustrate that the dynamics of mucoricin release during neutrophil-Mucorales interplay largely determine the infection outcome.

Finally, we evaluated cytokine immunotherapy as a therapeutic strategy of harnessing of neutrophil swarming by preventing mucoricin-induced apoptosis of neutrophils. Among the cytokines tested, we found that GM-CSF, a well-known cytokine with pleiotropic effects on the immune system, stabilized neutrophil clusters around germinating hyphae of Mucorales, enhanced ROS-mediated fungal killing, and prevented fulminant disease in vivo. Notably, a predominant mechanism of swarming enhancement by GM-CSF was the inhibition of mucoricin-induced apoptosis of neutrophils. The anti-apoptotic properties of GM-CSF against neutrophils are mediated via stabilization of Mcl-1^47^, an anti-apoptotic protein of the BCL-2 family. Collectively, our work paves the way for the design of novel host directed therapies that enhance neutrophil survival as a therapeutic strategy for mucormycosis.

Overall, our study shed light on early molecular events in host-Mucorales interplay and identified a new pathogenetic model of mucormycosis. Specifically, we discovered a novel mechanism of immunometabolic cross-talk between host metabolic factors regulating Mucorales pathogenicity and the efficiency of neutrophil swarming (**Graphical Abstract**). Under normal conditions, β-glucan exposure on germinating Mucorales triggers neutrophil swarming and results in fungal clearance. Furthermore, inhibition of fungal growth and mucoricin production by physiological albumin-bound FFAs facilitates neutrophil cluster stabilization and enhances fungal killing. DKA, hypoalbuminemia, or other mucormycosis-promoting conditions result in (a) delayed neutrophil recruitment and/or clustering, or (b) promote fungal growth and mucoricin expression, which collectively facilitate swarming disruption and invasive fungal disease. In contrast, GM-CSF and possibly other cytokines exert anti-apoptotic properties on neutrophils that enhance neutrophil swarming and prevent the development of mucormycosis. Collectively, our work provides novel perspectives in pathogenesis and management of an emerging, life-threatening disease with limited therapeutic options.

## Materials and Methods

### Microorganisms and culture conditions

*Aspergillus fumigatus* ATCC46645^32^, *Rhizopus delemar* (99–880)^6^, *R. arrhizus* (557969)^64^, *Cunninghamella bertholletiae* (506313)^64^, *R. delemar* RNA interference (RNAi) strains targeting mucoricin expression^6^, and *Mucor circinelloides* M16 strain^6^ have been previously described. *Rhizopus* spp, *C. bertholletiae,* and *A. fumigatus* were cultured on yeast extract agar glucose (YAG) plates at 37°C, for 5 days (d). *R. delemar* RNAi strains were grown on a chemically defined synthetic medium (per liter: 20g agar, 1.7g yeast nitrogen base (YNB) without amino-acids (Difco, #291920), 20g dextrose and 0.77g complete supplement mixture without uracil (CSM-URA, MP Biomedicals, #4511-212)). Furthermore, *M. circinelloides* M16 strain were cultured on Yeast Extract-Peptone-Dextrose (YPD) medium (per liter: 10g yeast extract, 20g peptone, 20g dextrose, 20g agar), supplemented with 0.2 mg/ml uracil and 0.2 mg/ml leucine. To generate *R. delemar* strain 99-880 spores lacking melanin, the fungus was grown on synthetic defined (SD) complete medium under two conditions: standard SD complete plates and SD complete plates supplemented with 1 mM of the copper chelator bathocuproinedisulfonic acid (BCS) for 5 d. Sporulation on BCS-containing plates induced copper deprivation, resulting in pigmentless spores, hereafter referred to as “albino” spores in subsequent experiments.

Fungal spores were harvested by using a sterile spreader in phosphate-buffered saline (PBS), filtered through 40 μm cell strainer (Falcon), washed twice with PBS following centrifugation at 3000 rpm for 5 minutes (min), and counted with a hemocytometer. When indicated, *Rhizopus* spores were inactivated by incubation with 4% paraformaldehyde (PFA) in PBS for 2 h at room temperature (RT).

Synchronization of *Mucorales* spore swelling was achieved by incubating 10^6^spores/mL at 28 °C in RPMI-MOPS supplemented with 2% glucose in Petri dishes. To suppress mucoricin expression during spore swelling, the culture medium was supplemented with 4.5 g/dL bovine serum albumin (BSA). The medium was then centrifuged using Amicon 3 kDa MWCO ultracentrifugal filters (Merck, #UFC5050) to remove albumin, generating an albumin filtrate containing albumin-bound free fatty acids (hereafter referred to as albumin-bound FFAs), as previously described^8^ [8]. This albumin filtrate was subsequently used to generate swollen *R. delemar* spores over ∼7 h at 28 °C.

For fluorescence labeling of fungal spores, 10^7^/ml dormant spores were stained over-night (O/N) at 4°C on a benchtop rotator either with 30 μg/ml Fluorescent Brightener 28 (CFW, Sigma-Aldrich, #475300) or with 200 μg/mL Fluorescein isothiocyanate isomer I (FITC Sigma Aldrich, #F7250)^65^ in 0.1 M NaHCO_3_. Afterwards, spores were washed three times with PBS and their concentration was adjusted according to experimental needs. For olorofim treatment, *Aspergillus fumigatus* dormant conidia were cultured in RPMI-MOPS medium supplemented with 0.1 mg/mL olorofim for 5 d at 28 °C.

### Preparation of *R. delemar* melanin ghosts

The isolation of melanin from *R. delemar* spores was performed as previously described^32^. Briefly, spores were treated with glycohydrolytic (Glucanex, Novozymes, Sigma-Aldrich #L1412) and proteolytic (proteinase K; Sigma Aldrich, #P2308) enzymes, guanidine thiocyanate (PanReac AppliChem, #A1107), and 6 M hydrogen chloride (HCl) pre-warmed at 100°C. This procedure led to electron-dense, spore shaped structures devoid of any underlying components, hereafter referred to as melanin “ghosts”.

### β-glucan and protein removal from *R. delemar* spores

The polysaccharide layer was enzymatically removed from swollen *R. delemar* spores. First, the spores were washed thrice with PBS, followed by a final wash in aqueous solution of1 M sorbitol and 0.1 M sodium citrate. Consecutively, the spores were incubated with 10 mg/ml β-1-3-D-glucanase (MegazymeE-LAMSE, SKU: 700004226) at 30°C O/N and repeatedly washed with PBS. Protein removal from the fungal cell wall of swollen spores was accomplished both enzymatically and chemically. For the enzymatic removal of surface proteins, spores were first treated with 4 M guanidine thiocyanate (AppliChem, A1107) at 4°C O/N and then with Proteinase K (Sigma-Aldrich, #P2308) in reaction buffer (10 mM Tris, 1mM CaCl_2_ and 0.5% SDS, pH 7.8) at 37°C, O/N. Afterwards, spores were washed three times with PBS. For the chemical removal of surface proteins, *R. delemar* spores were incubated with 1 M NaOH at 65°C for 1 h, centrifuged at 3.000 rpm for 10 min and re-treated with fresh 1 M NaOH before final washes with PBS.

### Human neutrophil isolation

Peripheral blood samples from healthy donors were obtained through the Blood Bank of University General Hospital of Heraklion. Each healthy volunteer provided written informed consent in accordance with the Declaration of Helsinki. Human peripheral blood neutrophils were isolated with double gradient of pancoll density(Pancoll human for granulocytes 1.119 g/ml (PANBIOTECH P04-60110); Pancoll human 1077 g/ml PANBIOTECH P04-60100) centrifugation technique as previously described^66^. Cell viability was assessed at 99% by trypan blue dye exclusion. Neutrophils were resuspended in RPMI medium (RPMI 1640, Gibco 11835-030) supplemented with 2% volume per volume (v/v) active human serum.

### Endothelial cells collection

Endothelial cells were collected from umbilical vein endothelial cells by the method of Jaffe et al^67^. The cells were harvested by using collagenase and were grown in M-199 enriched with 10% fetal bovine serum, 10% defined bovine calf serum, L-glutamine, penicillin, and streptomycin. Second-passage cells were grown to confluency in 24- or 96-well tissue culture plates (Costar, Van Nuys, CA) on fibronectin (BD Biosciences). All incubations were in 5% CO_2_ at 37°C. The reagents were tested for endotoxin using a chromogenic limulus amebocyte lysate assay (BioWhittaker, Inc.), and the endotoxin concentrations were less than 0.01 IU/ml. Endothelial cell collection was approved by Institutional Review Board at Los Angeles Biomedical Research Institute at Harbor-UCLA Medical Center.

### Human clinical samples

FFPE biopsies from patients diagnosed with mucormycosis were obtained from the Pathology Department, School of Medicine, at the University of Crete. Approval for the collection of all human samples used in the study was obtained from the Ethics Committee of the University of Heraklion, Crete, Greece (5159/2014, 10925/201 and 13-04-22/7970).

### Murine PMN isolation

Murine PMNs were isolated using a Percoll (Sigma-Aldrich P1644) triple gradient. The bone marrow was flushed out of tibia and femur using PBS (Thermo Fisher Scientific, #70011044) – 15Mm EDTA (Thermo Fisher Scientific, #15575020). Cells were centrifuged for 10 min at 300g and the pellet was resuspended in the same buffer. Neutrophils were purified by centrifugation for 30 min at 1,200 g -with no brakes, on a discontinuous gradient 52% (v/v), 69% (v/v) and 75% (v/v) Percoll in 0.15 M NaCl. After the recovery of cells from interphase between 69% and 75% Percoll, the cells were resuspended in a PBS-15 mM EDTA-1% BSA buffer and centrifuged for 10 min at 300g. The remaining erythrocytes were lysed using Endotoxin-Free Ultra-Pure water (Merck, #900000240432 TMS) and 0.3 M NaCl. Then, neutrophils were washed once with 2 ml HEPES buffer (2mM CaCl_2_, 5mM KCl, 1mM MgCl_2_, 140mM NaCl, 10mM HEPES), before finally resuspended in RPMI without phenol red, supplemented with 10% heat-inactivated serum. Cell viability was assessed at 99% by trypan blue dye exclusion. The purity of PMNs (identified as CD11b+/Ly6G+ cells) was >90% by flow cytometry.

### Animal Studies

Male and female *gp91^phox-/-^* (*Cybb^tm1Din^*) and control C57BL/6 (B6) mice (8- to 12-week-old) were used in all experiments. Mice were obtained from The Jackson Laboratory and maintained at the Specific pathogen free (SPF) facility of the Institute of Molecular Biology and Biotechnology (IMBB) in Heraklion Crete, Greece. The mice were maintained in grouped cages in a high-efficiency particulate air-filtered environmentally controlled virus-free facility (24 °C, 12/12h light/dark cycle), and fed a standard chow diet and water ad libitum. All experiments were approved by the local ethics committee of the University of Crete, School of Medicine, Greece, in line with the corresponding National and European Union legislation (animal protocols 17/07/2017-147075 and 22/03/2023-90477).

### Mouse model of pulmonary mucormycosis

In virulence studies, 8- to 12-week-old C57BL/6 (B6) mice were intratracheally infected with 5×10^6^ dormant *A. fumigatus* or *R. delemar,* or 2×10^6^ swollen *R. delemar* spores, as previously described^8,13^. For immunological studies, 8- to 12-week-old control C57BL/6 (B6) or *gp91^phox-/-^* mice were intratracheally infected with 5×10^6^ dormant *R. delemar* spores, euthanized at the indicated time point and BAL fluid was obtained.

The DKA model of mucormycosis in mice was performed as previously described^68^. In brief, freshly prepared streptozotocin (MedChem, HY-13753) (dissolved in ice-cold citrate buffer, pH 4.2, filter sterilized) was immediately administered to mice by intraperitoneal injection at a dose of 210 mg/kg. Ten days after streptozotocin injection, the mice were infected intratracheally with 5×10^6^ dormant *R. delemar* spores. Additionally, cortisone acetate (MedChem, MCE-HY17461A.50G) (dissolved in 0.1% Tween 80) was administered intraperitoneally to the mice (250 mg/kg) on days −2 and +3, relative to infection. For experiments assessing the potential of pro-survival benefit of GM-CSF (Peprotech, 315-03), mice received intraperitoneal doses of 5mg/kg recombinant GM-CSF, starting 6 days prior to infection with 2×10^6^ swollen *R. delemar* spores and occurring until day 3 post infection. In all experiments, after infection, two mice from each group were sacrificed for inoculum verification.

### Histopathological and immunohistochemistry/ immunofluorescence studies

Mice were euthanized, and their lungs were excised and fixed with 10% formalin before being embedded in paraffin and cut into 4-5 µm sections. Lung tissue sections were deparaffinized in xylene and rehydrated via an ethanol gradient (100%-70%). Hematoxylin and eosin (H&E) and MPO-PAS staining were performed according to the manufacturers’ instructions.

For neutrophil immunostaining in the lung, heat-induced antigen retrieval was performed to deparaffinize and rehydrate the lung sections through incubation in sodium citrate buffer (10 mM, 0.05% Tween 20, pH 6.0) for 40 min at 90–95°C, using a steamer (Philips). The tissue sections were allowed to cool for 30 min, washed three times with PBS and permeabilized with 0.2% gelatin/Triton X-100 0.25%/PBS for 15 min at RT, followed by blocking in 5% BSA/5% normal goat serum (Capricorn Catalog, #GOA1-A) (NGS)/PBS for 1 h at RT. The tissue slices sections were subsequently incubated at 4°C O/N with an anti-Ly6G (1:1000, Bio X Cell, #BE0075) antibody in 1% BSA/1% NGS/PBS. Washing 3 times with PBS was followed by incubation with goat anti-rat CF555 (1:500, Biotium, #) for 1 h, RT. For nuclear staining, the tissue sections were incubated with the nuclear dye TO-PRO-3 (1:2000, 642/661; Invitrogen), for 10 min at RT washed three times with PBS and quenched for autofluorescence by incubation in a solution of 10 mM CuSO_4_/50 mM NH_4_Cl for 5 min at RT. The same protocol was followed for mucoricin immunostaining in human sections.

For NETosis immunostaining in the lung, heat-induced antigen retrieval was performed to deparaffinize and rehydrate the lung sections through incubation in sodium citrate buffer (10 mM, 0.05% Tween 20, pH 6.0) for 40 min at 90–95°C, using a steamer (Philips). The tissue sections were allowed to cool for 30 min at RT, washed three times in PBS followed by blocking in 5% horse serum/ 0,05% Triton X-100/ PBS for 1 hour RT. The tissue slices were subsequently incubated at 4°C O/N with an anti-MPO (1:150, R&D systems, #AF3667) and an anti-CitH3 (1:500, Abcam, #ab281584) antibody in blocking buffer. Washing 3 times with PBS was followed by incubation with donkey anti-rabbit CF555 and donkey anti-goat (1:500, Biotium #20127, #20038) 1 hour RT. The tissue sections were incubated for 10 min at RT with the nuclear dye Ηoechst (1:2000, Thermo Fisher Scientific) washed thrice with PBS and quenched for autofluorescence by incubation in a solution of 10 mM CuSO_4_/50 mM NH_4_Cl. The same protocol was used for the human tissue sections.

For β-glucan and mucoricin immunostaining of *R. delemar*, the fungal spores were cultured at 37°C in RPMI-MOPS supplemented with 0.2% glucose. Spores at different stages of growth were fixed in 4% PFA for 1 h, followed by permeabilization in 0.1% Triton X-100 in PBS for 10 min, RT. The permeabilized spores were blocked with 1% NGS in PBS for 30 min, RT and subsequently incubated with the anti-Dectin1 antibody (0.15 mg/ml) for 1 h,RT. The spores were washed thrice with PBS and incubated with goat anti-mouse CF488A (Biotium) diluted 1:500 in PBS for 1 h RT. After washing with PBS, the spores were further blocked with 10% NGS in PBS for 1 h at RT and subsequently incubated with the anti-mucoricin antibody (1 mg/mL) for 2 h at RT. The spores were washed with Tris-buffered saline (0.01 M Tris HCl/0.15 M NaCl, pH 7.4) containing 0.05% Tween 20 and incubated with goat anti-rabbit CF555A (Biotium) diluted 1:500 in PBS for 1 h at RT.

### BAL staining

BAL cells were isolated from infected mice at the indicated time points and resuspended in RPMI supplemented with 1% heat-inactivated FBS and 1μΜ DRAQ5 (Biolegend, # 424101). The cells were seeded in 8 or 18-well iBidi μ-slides for 15 minutes, followed by blocking with anti-mouse CD16/32Trustain monoclonal antibody (0.5 mg/ml; TruStain FcX PLUS, Biolegend, # 156603) for 10 min. Then, the conjugated antibodies were added: anti-CD11c (1:200), anti-SiglecF (1:200), anti-Ly6G (1:100) together with 20 μΜ CFW for 1h RT. After extensive washes with PBS, the samples were observed before or after fixation with 4% PFA.

### Protocol for assessment of in vivo killing of *R. delemar* in BAL samples

BAL fluid was isolated at different time points following infection with FITC-labelled dormant *R. delemar* spores. Following centrifugation at 300 × g, BAL cells were resuspended in PBS and seeded at a concentration of10⁵ cells per well in a 96-well-plate. To discriminate extracellular from intracellular spores, 10 μg/ml CFW, a non-cell-permeable dye, was added to each well for 20 minutes in PBS. After six PBS washes to remove unbound dye, cells were lysed by sonication (40 Hz for 4 sec) and. The releases spores were incubated in RPMI without phenol red at 37 °C for 5 h to permit germination of both intracellular (FITC⁺, CFW⁻) and extracellular (FITC⁺, CFW⁺) fungal cells. Spore viability was determined by confocal microscopy by quantifying the proportion of germinated (viable) spores among the total counted.

For in vivo assessment of fungal killing in BAL samples, 8- to 12-week-old C57BL/6 (B6) mice or gp91^phox+/+^ and gp91^phox-/-^ mice were challenged by intratracheal infection with 5 × 10⁶ FITC-labelled dormant *R. delemar* spores. BAL fluid was collected at various time points post infection (0, 3, 6, 12, 24, and 48 h). At each time point, two mice per group were euthanized via ketamine/xylazine overdose. BAL samples were processed as described above.

### Measurement of Reactive Oxygen Species (ROS) production

ROS production was quantified by means of a dichlorofluorescein assay. The stock solution of dichlorofluorescein diacetate (DCFH-DA) (Thermo Fisher Scientific, Catalog #D399) was dissolved prepared in dimethyl sulfoxide (DMSO) at a final concentration of 10 mM. The isolated neutrophils were seeded in 18-well plates in RPMI supplemented with 10% heat-inactivated FBS. DCFH-DA was added 30 min prior to infection at final concentration of 10 μM at 37°C. At the indicated time point of infection, the fluorescence intensity of DCFH-DA, corresponding to intracellular ROS levels, was assessed in individual cells by confocal microscopy upon excitation with a 488 nm laser line and detection through a 521/38 nm bandpass emission filter.

### Blocking of β-glucan receptors in neutrophils

For neutralization of Dectin-1 and Complement Receptor 3 (CR3), isolated neutrophils were seeded for 15 minutes at 37 °C, and then treated with 10μg/ml the anti-Dectin-1 (InvivoGen, Catalog #mabg-mdect-2) for 1 h or 20μg/ml anti-CD11b (M1/70, Thermo Fisher Scientific, Catalog #14-0112-82) for 30 min along with their isotype controls IgG2a (Bio X Cell, Catalog #BE0089) and IgG2b (R&D Systems, Catalog #MAB0061). Following antibody incubations, β-glucan particles labeled either with FITC or CFW, were added to the neutrophils at a 20:1 cell-to-particle ratio (1 × 10⁵ cells: 5 × 10³ particles) and incubated for 30 min at 37°C. Additionally, *R. delemar* spores were introduced in the presence of 1ng/ml GM-CSF and incubated for up to 5 h for *R. delemar* growth inhibition assessment. Following incubations, cells were fixed with 4% PFA for 10 min, RT.

### Annexin V/PI assay

To evaluate the cell death of neutrophils 15 and 30 min of infection, bone-marrow isolated neutrophils in RPMI suspension supplemented with 10% heat-inactivated FBS were seeded in 18-well plates and challenged with either swollen control (empty RNAi) or mucoricin RNAi *R. delemar* spores. At each time points, the Annexin V/PI staining was performed according to the manufacturers’ instructions.

### Apoptosis inhibition assay

For apoptosis inhibition assay, bone-marrow isolated neutrophils were seeded in 18-well plate for 15 min at 37°C. Then, GM-CSF, z-VAD-FMK (MedChem, MCE-HY16658B), z-DEVD-FMK (MedChem, MCE-HY12466.1), were added for 1 h. Then, cells were challenged with swollen *R. delemar* spores for 3 h, before acquisition.

### Recombinant mucoricin production

Heterologous expression of mucoricin gene in *E. coli* was performed by extracting total RNA from *R. delemar* 99-880 hyphae grown on YPD broth for overnight and reverse transcribed into cDNA using the High-Capacity cDNA Reverse Transcription Kit (Fisher Scientific, Cat # 43-688-14). The entire ORF of mucoricin was PCR amplified from cDNA by Phusion high-fidelity PCR Kit (New England Biolabs) using primers 5’-CGCGGATCCATGTATTTCGAAGAAGGCCGC-3’ and 5’-CCGGAATTCTTATCCTTCAAATGGCACTAATTCCCA-3’. The amplified PCR product was cloned into pGEX-2T *E. coli* BL21 expression vector in the BamHI and EcoRI sites downstream of the Tac promoter. The generated *E. coli* expression vector was transformed into *E. coli* BL21 and transformants were screened on LB broth with ampicillin. The produced recombinant mucoricin should be GST-tagged on the N-terminus. *E. coli* cells transformed with empty plasmid served as negative control. The expression of mucoricin was induced by 0.2 mM of isopropyl β-D-thiogalactoside (IPTG) in LB broth containing 100 µg/ml ampicillin and purified using glutathione sepharose 4B resin according to the manufacturer’s protocol (Thermo Scientific). Protein purity (>95%) was confirmed by SDS-PAGE and Coomassie staining. Endotoxin levels were (below 0.1 EU/ml where measured). Purified Mucoricin was concentrated to 1-2 mg/mL, aliquoted, flash-frozen, and stored at −80 °C. Protein identity was validated by LC-MS and activity was confirmed using ribotoxin activity assay.

### Mucoricin binding on β-glucan assay

To accomplish binding of mucoricin on β-glucan particles, 100 μg of purified mucoricin was added to 10^6^/ml β-glucan particles in ultra-pure water. The mixture was incubated at 37°C in a bench-top rotator for 3 h. After the incubation, the particles were washed with PBS and part of them were stained for mucoricin, while others were used for swarming assays.

### Assesment of neutrophil cluster disruption by released mucoricin

To evaluate the effect of mucoricin released during fungal growth on neutrophil cluster disruption, bone marrow–derived neutrophils were stimulated with β-glucan particles in the presence or absence of purified recombinant mucoricin, which was added to the culture medium at 30 min at a final concentration of 100 μg/mL. After 3 h of incubation, Sytox Green (1 μM) was added to each condition, and neutrophil cell death was immediately quantified by confocal microscopy.

### Cytotoxicity Assay

Mucoricin-induced cell death was additionally assessed by measuring the release of cytoplasmic lactate dehydrogenase (LDH) using the Promega CytoTox 96® Non-Radioactive Cytotoxicity Assay (#G1780) following the manufacturer’s instructions. LDH is rapidly released into the culture medium when the plasma membrane is compromised, which occurs during cell death. The assay quantifies LDH activity based on the enzymatic conversion of lactate to pyruvate, generating NADH, which in turn reduces a tetrazolium salt (INT) to form a red, water-soluble formazan dye. The amount of formazan formed is directly proportional to the number of dead or damaged cells.

A549 (human lung epithelial), HUVEC (human umbilical vein endothelial), and primary mouse neutrophils were used in this assay. A549 and HUVEC cells were seeded in 96-well plates at a density of 1 × 10⁵ cells/well and allowed to adhere overnight. Mouse neutrophils were freshly isolated from bone marrow using the MojoSort™ Neutrophil Isolation Kit (BioLegend, #480058) and plated immediately at the same density. All cell types were treated with r-Mucoricin at 50 µg/mL for 6 h at 37°C in a humidified 5% CO₂ incubator. Untreated cells served as negative controls, while total cell death was induced in parallel wells using lysis buffer provided in the kit to determine maximum LDH release. After treatment, 25 µL of supernatant from each well was transferred to a new 96-well plate and combined with 25 µL LDH substrate solution. The reaction was incubated for 30 min at room temperature in the dark. Following incubation, 25 µL of Stop Solution was added, and absorbance was measured at λ = 490 nm (OD₄₉₀) using a microplate reader (BioTek).

### Neutrophil swarming under different pH conditions

To examine the effect of pH on neutrophil response, neutrophil swarming was evaluated following stimulation with β-glucan particles in RPMI media supplemented with 10% heat-inactivated FBS. RPMI was supplemented with 25mM MOPS and the pH of the medium was adjusted to 6.8, 6.9, 7.0, 7.2 and 7.4. The neutrophils were incubated at 37°C in the different pH conditions for 1h, before being challenged with β-glucan particles.

### BHB-mediated swarming inhibition

To examine the effect of BHB on neutrophil response, neutrophil swarming was evaluated using β-glucan particles in RPMI supplemented with 10% heat-inactivated FBS. RPMI was supplemented with 10mM BHB, as previously described^11^. The neutrophils were incubated at 37°C in the different pH conditions for 1h, before being challenged with β-glucan particles.

### Live imaging and acquisition

All images were acquired using a spinning disk confocal system (Dragonfly 200, Andor) mounted on a motorized inverted microscope (Nikon Eclipse Ti2-E). The system was equipped with an Andor Sona sCMOS 4.2B-6 (2048 × 2048 pixels) and an iXon EM-CCD (1024 × 1024 pixels) camera, four laser lines (405, 488, 561, and 633 nm), and bandpass emission filters at 445/46 nm, 521/38 nm, 594/43 nm, and 698/77 nm. Imaging was performed within a humidified incubation chamber (Tokai Hit Top Incubator) with the use of Sona sCMOS camera. The following objectives were used: 10×/0.45 (CFI Plan Apochromat Lambda, MRD00105), 20×/0.80 (CFI Plan Apochromat Lambda D 20X, MRD70270), 40×/1.15 water (CFI Apochromat LWD Lambda S 40XC WI, MRD77410), and 60×/1.4 oil (CFI Plan Apochromat Lambda, MRD). Nikon immersion oil type F2 was used when indicated. All images were obtained via z-stacks, deconvolved with Fusion software version 2.3.0.44 (Andor – Oxford Instruments), and further processed in Imaris 10.1 (Andor – Oxford Instruments) for contrast adjustment, area selection, color combining and scale bar addition. Certain images were acquired with a Leica TCS SP8 confocal microscope with a 63x lens and analyzed with the use of LASX and Fiji.

### Statistical analysis

Statistical analyses were performed using Prism (GraphPad Software). A p value of < 0.05 was considered statistically significant for all variables tested. The data were processed and visualized via GraphPad Prism (version 9.04). The statistical methods used to determine significance and p-values of each graph are provided in the figure legends. Neutrophil–fungal interactions were classified as follows: (a) **phagocytosis**, defined as the complete engulfment of a fungal spore; (b) **swarming**, defined in *in vivo* experiments as the formation of at least two neutrophil layers surrounding fungal spores, or ex vivo as a cluster comprising at least one layer of more than five neutrophils; and (c) **association**, defined as the adhesion of a neutrophil to a fungal spore. All experiments were performed in a minimum of three independent replicates, unless otherwise specified. For each replicate, at least 50 neutrophil–fungal interaction events were analyzed to assess swarming behavior.

### Financial support

S.B. was supported by the General Secretariat for Research and Technology (GSRT) and the Hellenic Foundation for Research and Innovation (H.F.R.I.) Under the “3^rd^ call of H.F.R.I. for PhD Candidates”. M.S. was supported by Fondation Santé. A.V. was supported by Fondation Santé. G.C. is supported by an ERC Consolidator Grant (iMAC-FUN, #864957), an Horizon 2020–Research and Innovation Framework Programme from Europe, H2020-SC1-BHC-2018-2020 (HDM-FUN #84750), and a “la Caixa” Foundation Health Research Grant (TRANS-CPA, #LCF/PR/HR17/52190003). G.C. and M.S. are supported by the General Secretariat for Research and Innovation of Greece Grant PRO-sCAP (Project Code TAEDR-0541976) carried out within the framework of the National Recovery and Resilience Plan Greece 2.0 and funded by the European Union-Next Generation EU. A.S.I., Y.G., and S.D., are supported by Public Health Service grant no R01 AI063503 and P01 AI186818-01.

## Supporting information

Supplementary Images 1-9

## Author Contributions

S.B established protocols, performed and analyzed most of the experiments, and participated in the writing of the manuscript; M.S. established protocols, performed experiments in mouse and human neutrophils and analyzed data; T.A. participated in the establishment of functional assays; A.V. analyzed data and performed experiments for establishment of in vivo killing in BAL; E.D. and A.K. provided human samples, performed histopathological studies, and analyzed data; Y.G. performed experiments on purification of mucoricin and analyzed data; S.D. performed experiments on mucoricin-mediated cell death in different cell types; A.S.I. participated in discussions and experimental design throughout the study, edited the manuscript, and provided resources; G.C. conceived and supervised the study, was involved in the design and evaluation of all of the experiments, and wrote the manuscript along with comments from the co-authors.

## Acknowledgments

The authors have no conflicting financial interests related to this work. We thank all members of the Chamilos Lab for their valuable discussions. The authors would like to thank Dr Konstantinos Kampas for assistance in establishment of human tissue protocols on immunostaining of neutrophils; the technical assistance of the perinatal nurses of the Harbor-UCLA General Clinical Research Center for collection of umbilical cords, the personnel of the Pathology facility at the University Hospital of Heraklion and Dr Maria Makkou for assistance with the identification and processing of tissue biopsy samples from patients with mucormycosis.

## Supplementary Figures

**Figure S1, related to** Fig. 1 **Ex vivo murine neutrophil swarming against towards Mucorales ex vivo. (A)** Data on quantification of the number of spores engaged in different neutrophil BAL cell responses in the lungs of immunocompetent mice infected with either *R. delemar* at 24 h. **(B)** Representative images of murine bone-marrow neutrophils infected either with dormant *R. delemar* spores or *A. fumigatus* conidia, 4 hours post-infection. Scale bar, 10 μm. **(C)** Representative images of murine bone-marrow neutrophils infected either with dormant *C. bertholletiae* or *M. circinelloides* spores, at 4 h of infection. Scale bar, 30 μm. **(D)** Data on the quantification of different neutrophil responses in murine bone-marrow neutrophils infected either with *C. bertholletiae* or *M. circinelloides*.

**Figure S2, related to Fig. 1. Assessment of in vivo killing of *R. delemar* in the lung. (A)** Schematic illustration of the protocol for assessment of in vivo killing of intra- and extracellular fungal spores. **(Β)** Percentage of killed *R. delemar* spores 3-, 6-, 12- and 24 h post infection of immunocompetent C57BL/6 (B6) mice with dormant fungal spores. The killing of intracellular spores begins earlier than the killing of the extracellular ones -3 and 6 h respectively, since there are many resident macrophages that phagocytose the spores, while neutrophils infiltrate the lung approximately 3 hours post infection. **(C)** Mice were sacrificed at the indicated time points, lungs were homogenized, and fungal loads were assessed by CFU plating. \*\*\**P* = 0.005, \*\*\*\**P* < 0.0001 One-way ANOVA. **(D)** Mean fungal area within macrophages in BAL of *gp91^phox +/+^* mice and *gp91^phox -/-^* mice, infected with dormant *R. delemar* spores (n= 2/group). *P*=0.7094, Unpaired t-test.

**Figure S3, related to** Figure 2**. Human neutrophils swarm towards swollen *R.delemar* ex vivo. (A)** Representative image of human neutrophils challenged with swollen *R. delemar* spores treated with PFA. Scale, 30 μm. **(B)** Data on quantification of the number of spores engaged in different neutrophil responses in the case of human neutrophils challenged with swollen *R. delemar* spores treated with PFA, 1 h post infection.

**Figure S4,** related to Figure 2. Neutrophil swarming is induced by a cell wall molecule exposed on germinating *R. delemar* spores after melanin removal*. (*A) Representative image of murine bone-marrow derived neutrophil challenged with melanin particles isolated form *R. delemar* spores. The melanin particles are completely inert to neutrophils. Scale bar, 30 μm. **(B)** Data on quantification of bone marrow-isolated neutrophils, challenged with melanin particles. \*\**P*=0.0048, Unpaired t-test. **(C)** Representative images of murine bone-marrow derived neutrophil challenged with PFA-killed albino spores and control ones. Scale bar, 30 μm.

**Figure S5, related to** Figure 2**. Olorofim-treated swollen *A.fumigatus* spores induce neutrophils swarming. (A)** Representative images of murine bone-marrow derived neutrophils infected with swollen *A. fumigatus* in the presence or absence of olorofim. Scale bar, 20 μm **(B)** Data on quantification of different neutrophil responses in the lungs of immunocompetent mice infected with *A. fumigatus* in the presence or absence of olorofim. (**C**) Representative images of immunofluorescence staining of β-glucan of olorofim-treated *A. fumigatus* conidia. Scale bar, 10 μm.

**Figure S6,** related to Figure 3. Exposure of a cell wall-associated virulence factor at ∼4 hours of *R. delemar* spore induces fulminant pneumonia. Survival rates of immunocompetent mice infected with dormant, 2- and 4 h swollen *R. delemar* spores (n=5/group). The nonparametric log-rank test was used to determine differences in survival rates.

**Figure S7, related to** Fig. 4**. Genetic deletion of mucoricin expression stabilizes neutrophil clusters around germinating Mucorales spores.** Representative time lapse images of mouse bone-marrow neutrophils challenged with either control RNAi or mucoricin RNAi *R. delemar strain*, or β-glucan. Scale bar, 30 μm.

**Figure S8, related to** Fig 4**. Both released and cell wall–bound mucoricin induces neutrophil apoptosis and swarming disruption. (A)** Quantification of cluster area of murine bone-marrow derived neutrophils challenged with swollen *R. delemar* spores, pre-exposed to 5 mM galactose for 1 h. *\*\*\* P =0.0001,* Mann-Whitney test. **(B)** Murine bone-marrow isolated neutrophils, infected with β-glucan particles, were challenged with 100 μg/ml purified mucoricin toxin. Scale bar, 20 μm. **(C)** Quantification of Sytox Green^+^ cells around β-glucan particles in the presence of purified mucoricin toxin. **** *P* <0.0001, Mann-Whitney test. **(D)** Comparative susceptibility of lung epithelial cells (A549), endothelial cells (HUVECs), or neutrophils to killing by recombinant (r) mucoricin following 6 h of incubation at a final concentration of 50 μg/mL. **** *P*< 0.0001, One-way ANOVA

**Figure S9, related to Fig. 4. Pre-exposure of swollen *R. delemar* spores to albumin-bound FFAs restores neutrophil swarming and prevents mucoricin-induced neutrophil death in vivo and ex vivo. (A)** Representative images of BAL isolated from infected mice with swollen *R. delemar* spores with or without pre-exposure to albumin-bound FFAs at 6 h of infection (n=2/group). Scale bar, 30 μm. **(B)** Quantification of dead cells in neutrophil clusters in BAL isolated from infected mice with swollen *R. delemar* spores with or without pre-exposure to albumin-bound FFAs at 6 h of infection. *\*\*\*P* < 0.0001, Unpaired t-test. **(C)** Quantification of number of cells per cluster in in BAL isolated from infected mice with swollen *R. delemar* spores with or without pre-exposure to albumin-bound FFAs at 6 h of infection. *\*\*\*P* < 0.0001, Unpaired t-test. **(D)** Representative images of murine bone-marrow isolated neutrophils, infected with swollen *R. delemar* spores with or without pre-exposure to albumin-bound FFAs ex vivo, at 1 h of infection. Scale bar, 30 μm. **(E)** Data on quantification of number of cells per cluster of murine bone-marrow isolated neutrophils, infected with swollen *R. delemar* spores with or without pre-exposure to albumin-bound FFAs ex vivo, at 1 h of infection. \*\*\**P*=0.0010, Unpaired t-test.

## Supplementary Videos

**Supplementary Video 1.** Time lapse movie on the kinetics of swarming induction in mouse bone marrow derived neutrophils infected with live *R. delemar* dormant spores, corresponding to Fig. 2A.

**Supplementary Video 2.** Time lapse movie on the dynamics of swarming formation in mouse bone marrow derived neutrophils infected with live swollen spores *of R. delemar* or *A. fumigatus*, corresponding to Fig. 3C.

**Supplementary Video 3.** Time lapse movie on the dynamics of swarming formation in mouse bone marrow derived neutrophils infected with live swollen spores of control RNAi or mucoricin RNAi *R. delemar* or large β-glucan particles, corresponding to Fig. 4A.

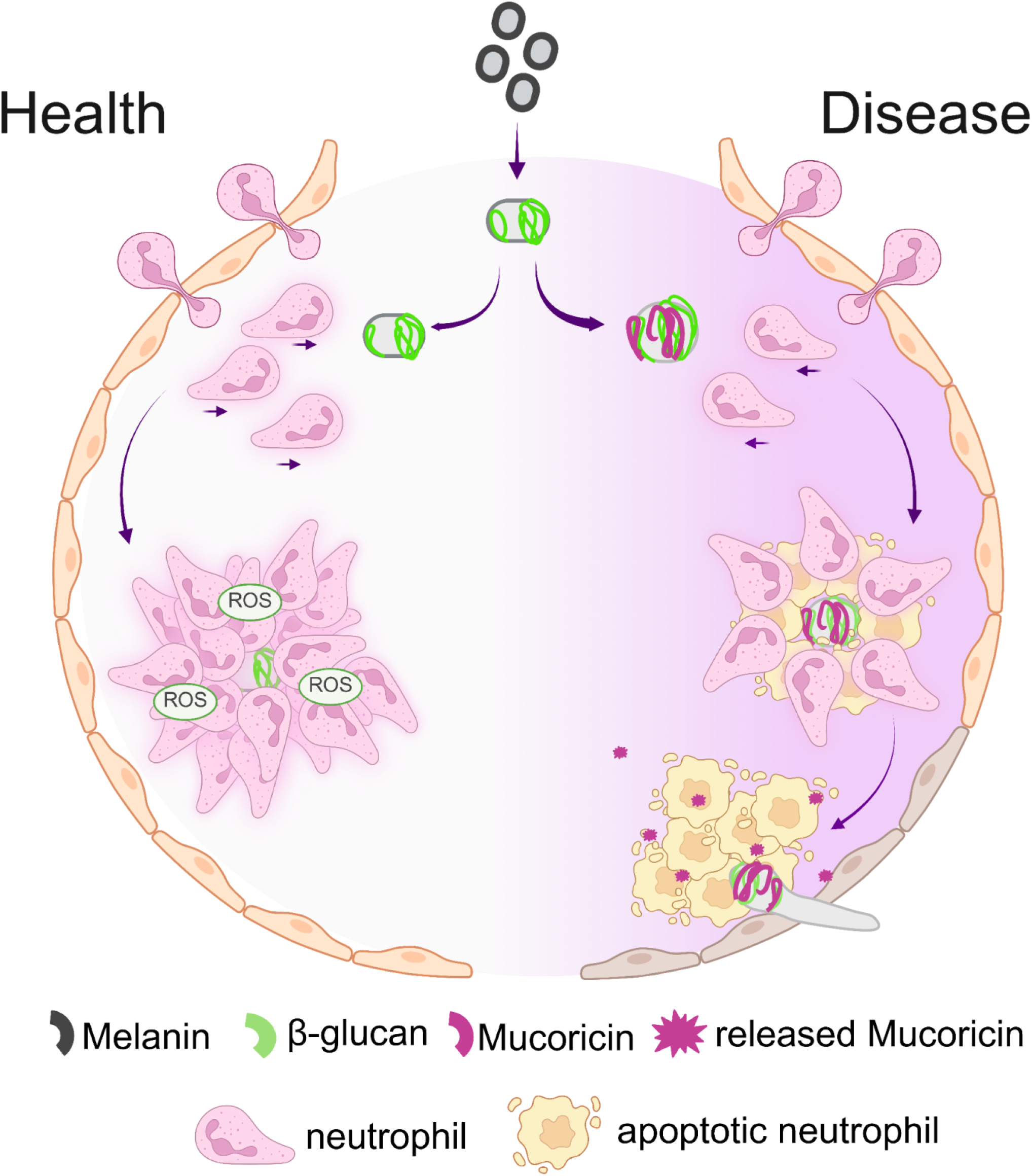

