## Supplementary Images 1-9 for "Mucoricin binding to β-glucan sites on germinating Mucorales spores disrupts neutrophil swarming to promote pathogenicity"

### Supplementary Figure 1

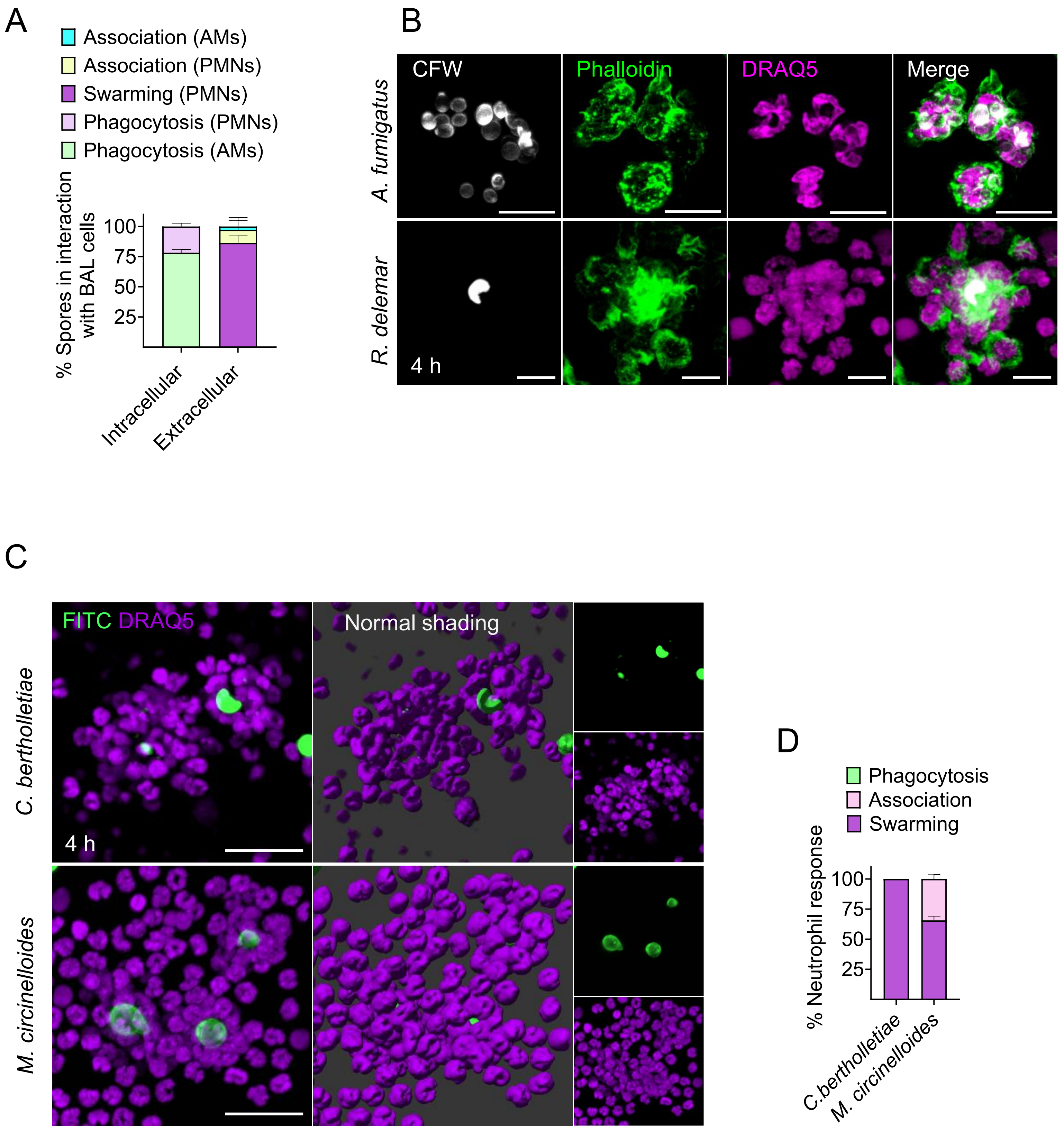

### Supplementary Figure 2

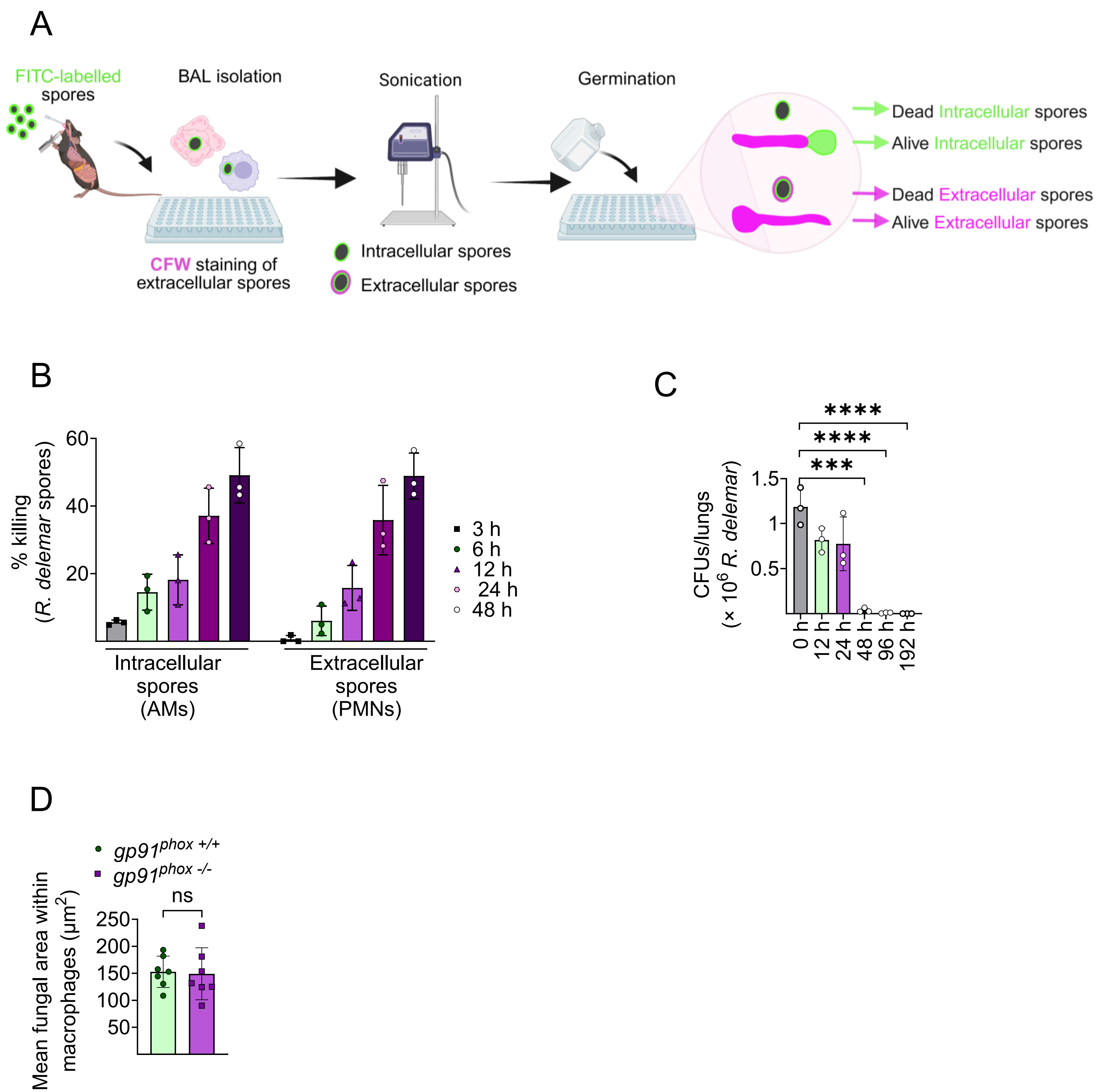

### Supplementary Figure 3

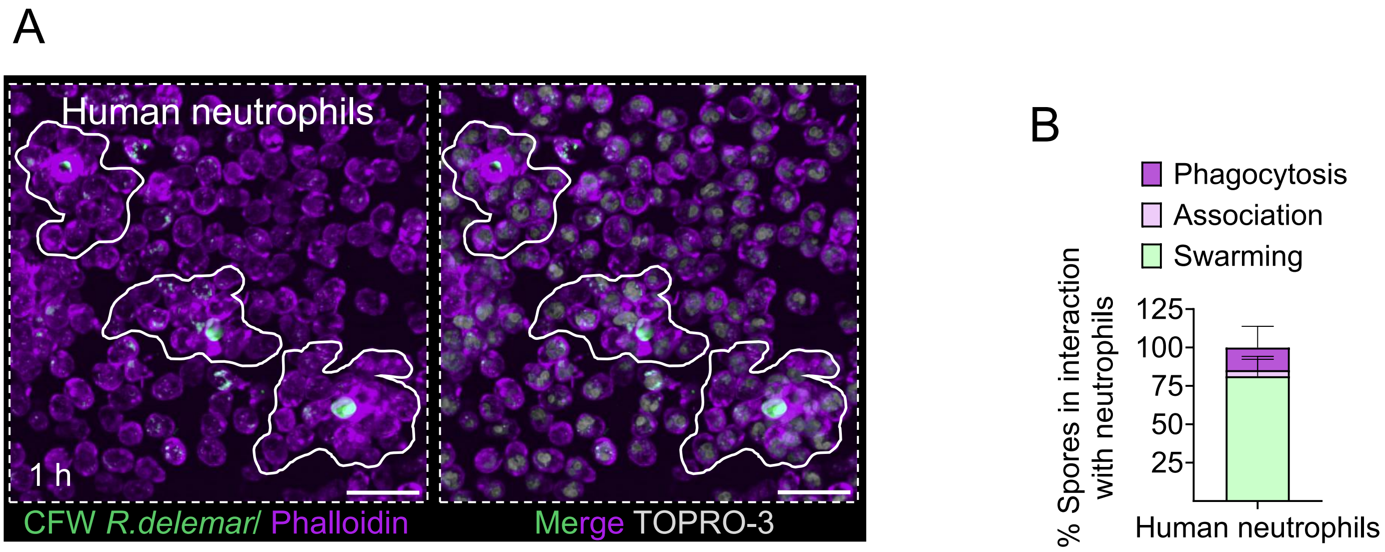

### Supplementary Figure 4

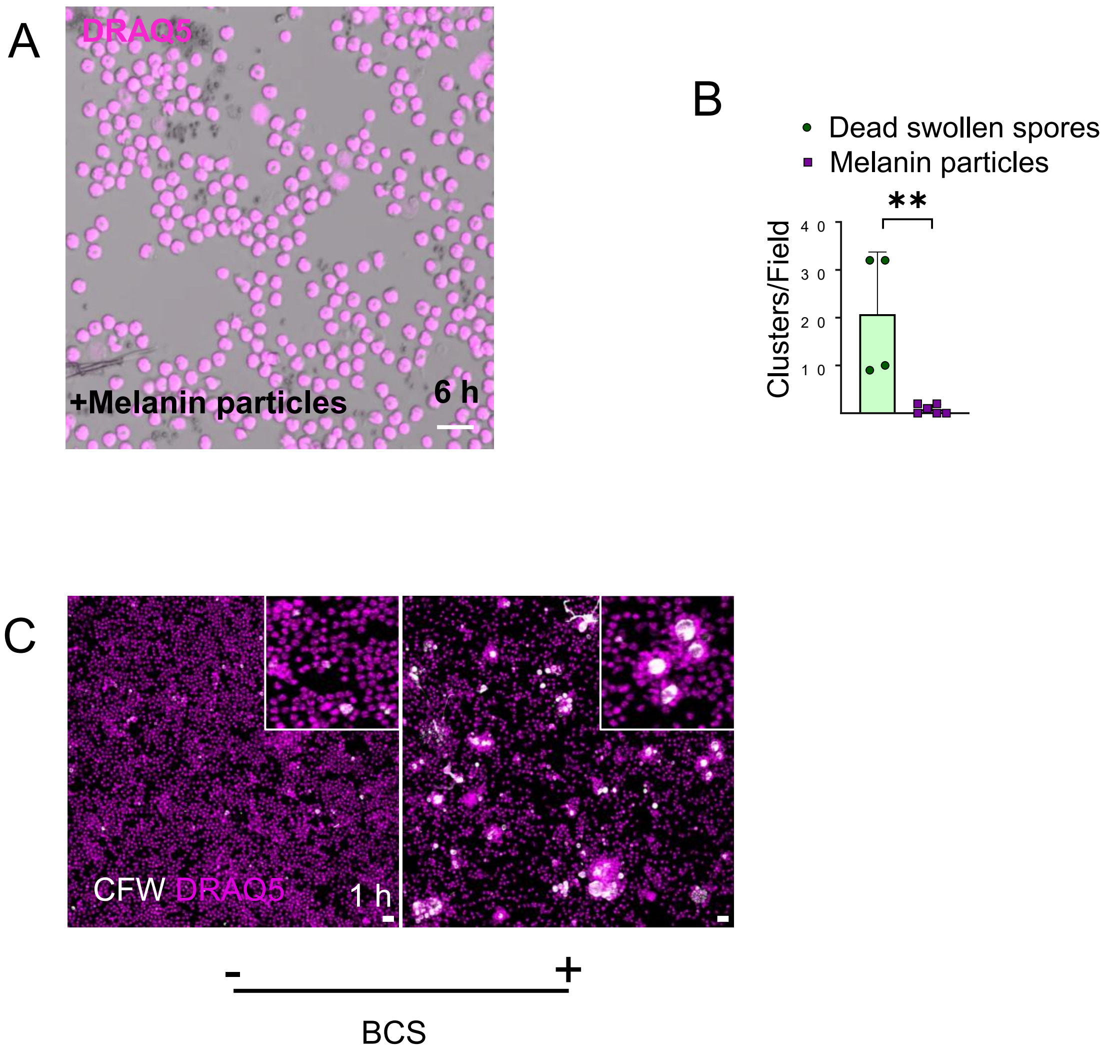

### Supplementary Figure 5

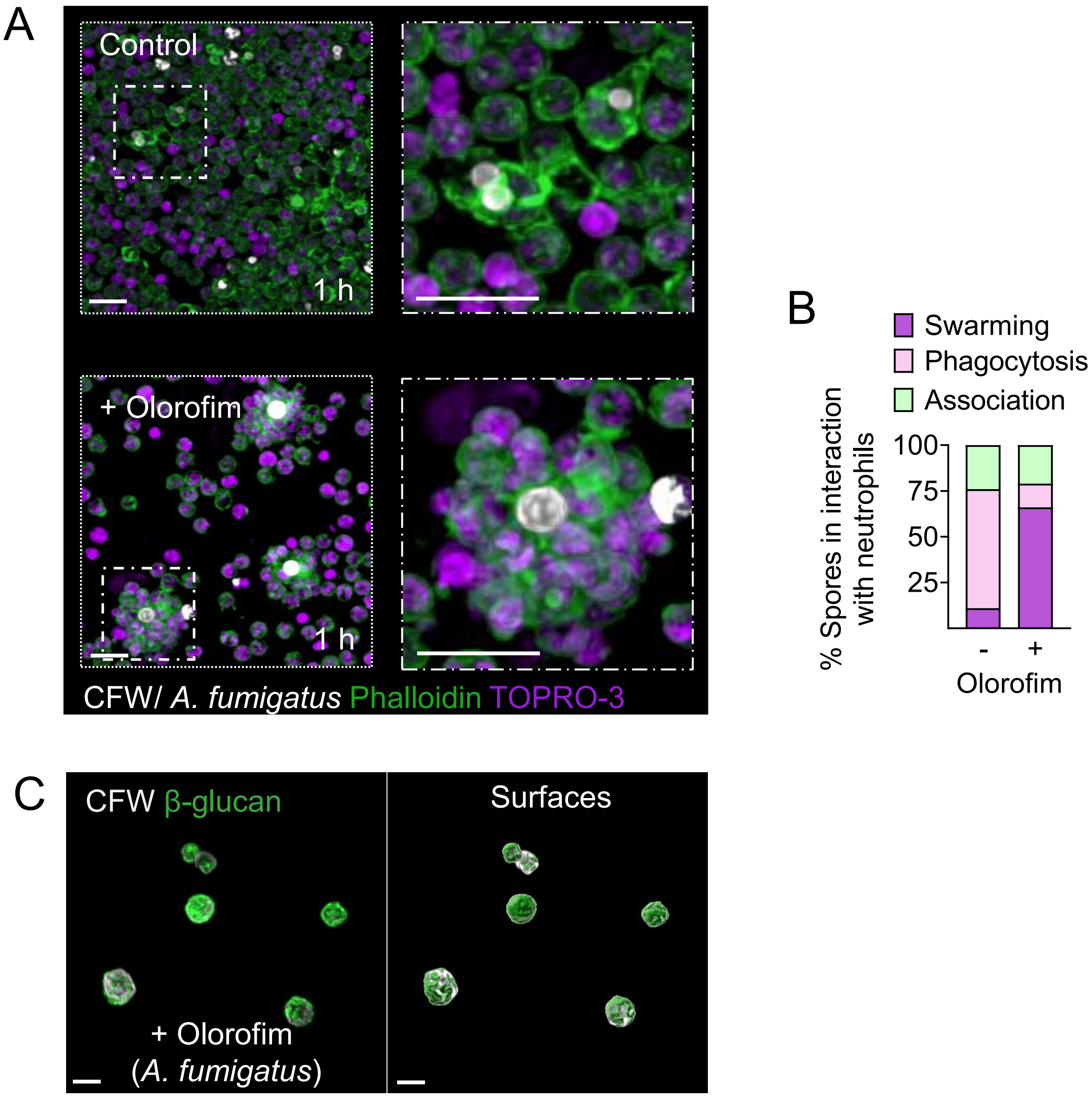

### Supplementary Figure 6

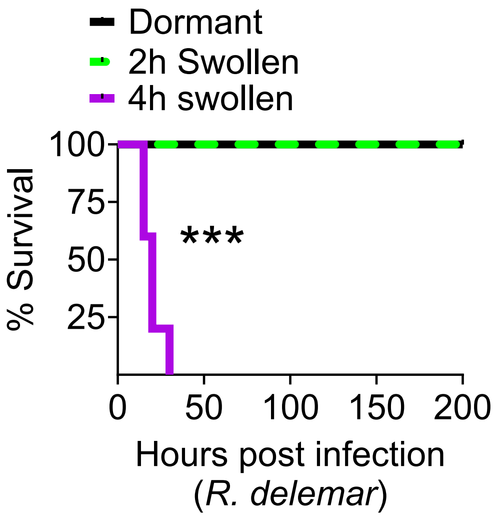

### Supplementary Figure 7

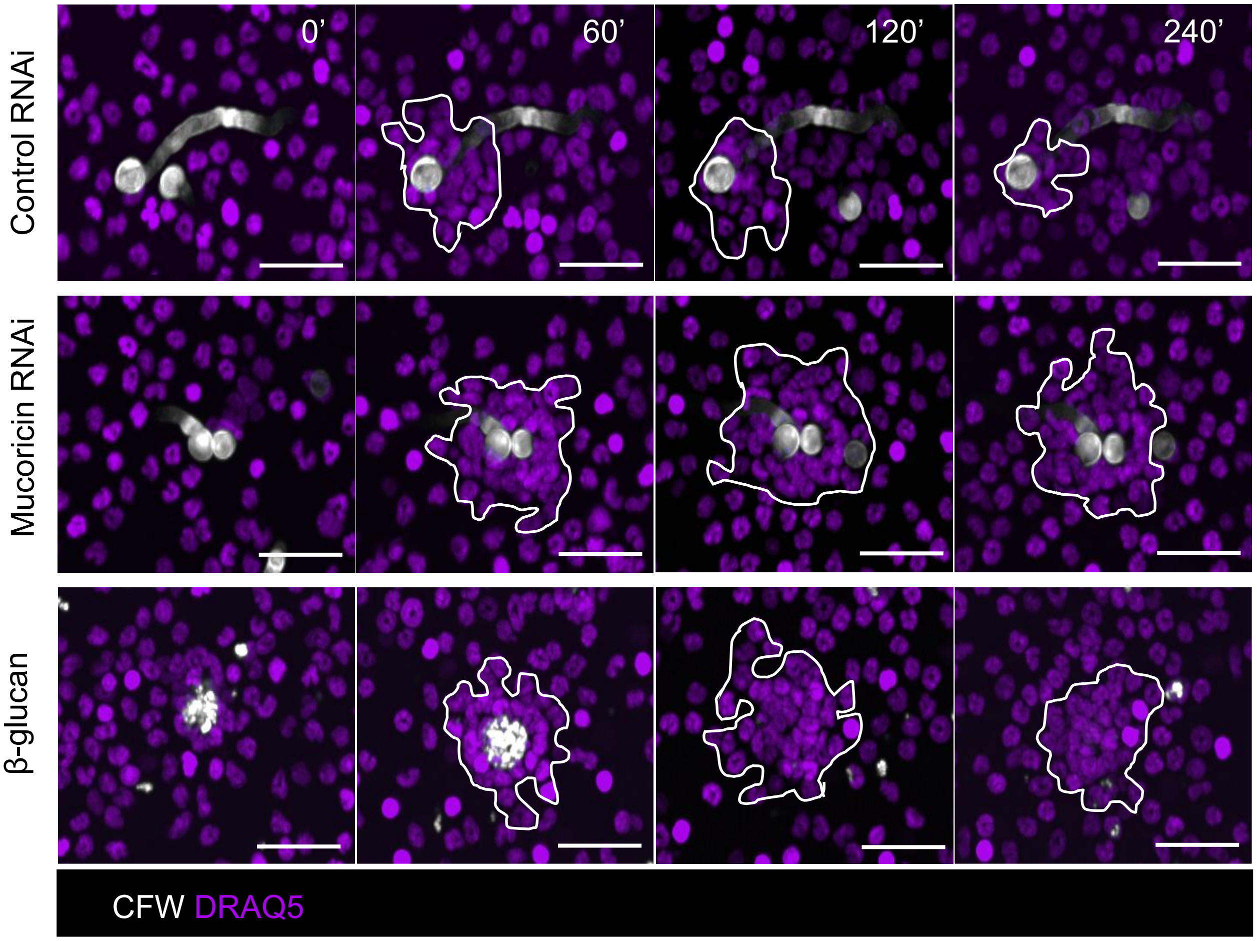

### Supplementary Figure 8

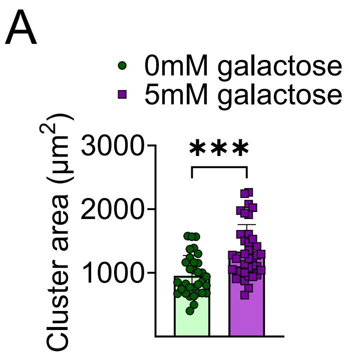


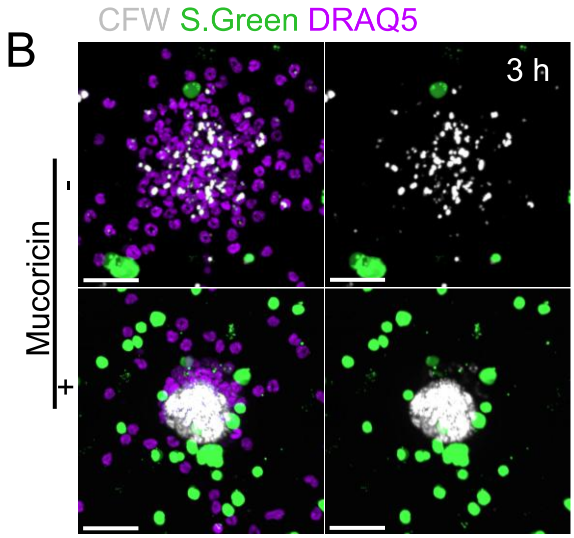


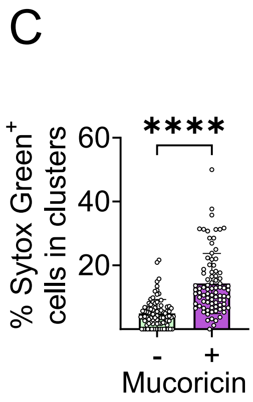


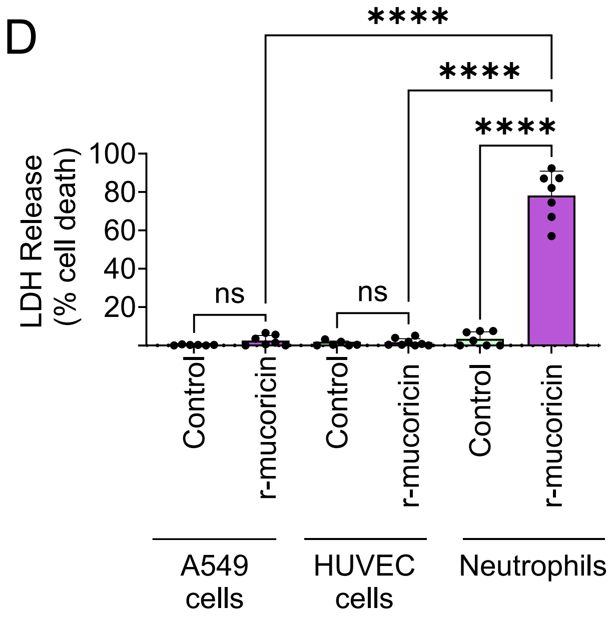

### Supplementary Figure 9

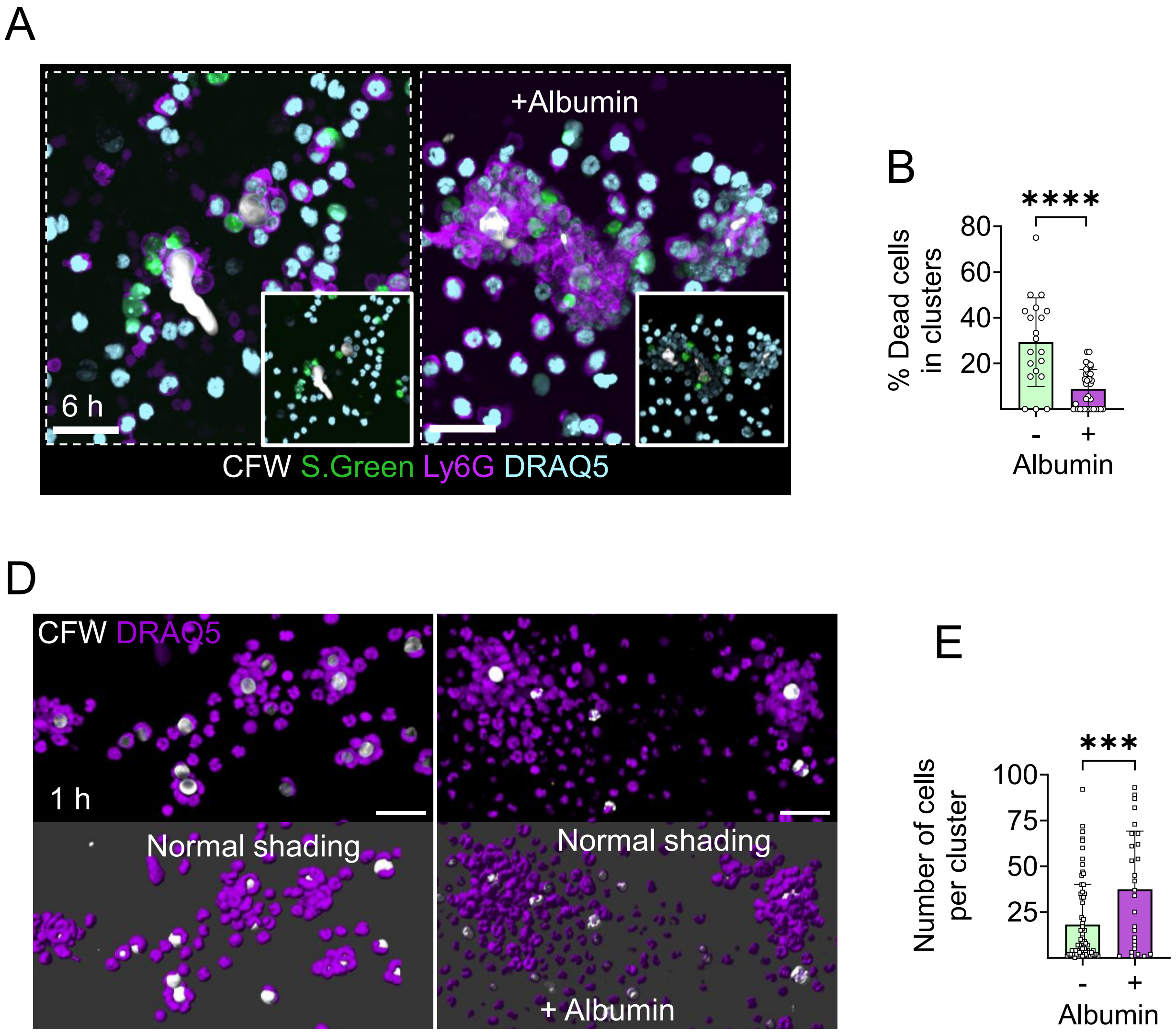
